# Freeform robotic optical coherence tomography beyond the optical field-of-view limit

**DOI:** 10.1101/2024.05.21.595073

**Authors:** Raymond Fang, Pengpeng Zhang, Tingwei Zhang, Daniel Kim, Edison Sun, Roman Kuranov, Junghun Kweon, Alex Huang, Hao F. Zhang

**Affiliations:** Department of Biomedical Engineering, Northwestern University, Evanston IL; Department of Mechanical Engineering, Northwestern University, Evanston IL; Department of Chemical Engineering, Northwestern University, Evanston IL; Department of Ophthalmology, University of California San Diego, San Diego CA; Opticent Health, Evanston IL

## Abstract

Imaging complex, non-planar anatomies with optical coherence tomography (OCT) is limited by the optical field of view (FOV) in a single volumetric acquisition. Combining linear mechanical translation with OCT extends the FOV but suffers from inflexibility in imaging non-planar anatomies. We report the freeform robotic OCT to fill this gap. To address challenges in volumetric reconstruction associated with the robotic movement accuracy being two orders of magnitudes worse than OCT imaging resolution, we developed a volumetric registration algorithm based on simultaneous localization and mapping (SLAM) to overcome this limitation. We imaged the entire aqueous humor outflow pathway, whose imaging has the potential to customize glaucoma surgeries but is typically constrained by the FOV, circumferentially in mice as a test. We acquired volumetric OCT data at different robotic poses and reconstructed the entire anterior segment of the eye. The reconstructed volumes showed heterogeneous Schlemm’s canal (SC) morphology in the reconstructed anterior segment and revealed a segmental nature in the circumferential distribution of collector channels (CC) with spatial features as small as a few micrometers.

## Introduction

Optical coherence tomography (OCT) has revolutionized fundamental investigations in various medical disciplines, including cardiology, dermatology, and ophthalmology^1-3^. In particular, OCT has transformed the clinical management for nearly all blinding diseases^4-7^. OCT is a non-invasive microscopic imaging modality that acquires volumetric data by detecting back-scattered photons at each optical illumination position and translating the optical focus of illumination to cover the region of interest (ROI). Such a data acquisition scheme has remained essentially unchanged since OCT was first reported in 1991.

All OCT systems have a finite imaging volume, defining the spatial coordinates that are observable by the design, which is related to the field of view (FOV) and the depth of the field (DOF). As illustrated in Fig. 1a, the optical implementation determines the FOV, and the DOF is governed by the optical penetration depth and spectral domain sampling density^8^. The limited OCT imaging volumes are often insufficient to cover large ROIs with spatial heterogeneity. Extending the imaging volume is achieved by implementing 3-degree-of-freedom (DoF) translational motions in the Cartesian coordinates (Fig. 1b), as seen in nearly all modern microscopes^9^. Although this strategy enlarges the compound imaging volume (CIV) for stationary, flat samples, it faces several challenges in imaging non-planar anatomical structures. Existing hybrid scanning only allows a fixed, predefined optical illumination direction. Lacking the capability to adjust the optical illumination direction, especially for samples with non-planar features, true for nearly most structures *in vivo*, results in nonconformal imaging, where the incident illumination is oblique to the normal vector of the tissue surface. This nonconformal imaging reduces imaging accuracy due to degraded signal-to-noise ratio (SNR) and en-face projection artifacts. Moreover, mechanical translation of imaging volumes for highly non-planar sample surfaces may result in unused data in its CIV, leading to unnecessarily long data acquisition and processing time, and large storage space. Hence, an ideal OCT should maximize the CIV, follow the sample contour for conformal scanning, and adjust the illumination direction as needed.

**Fig. 1.**
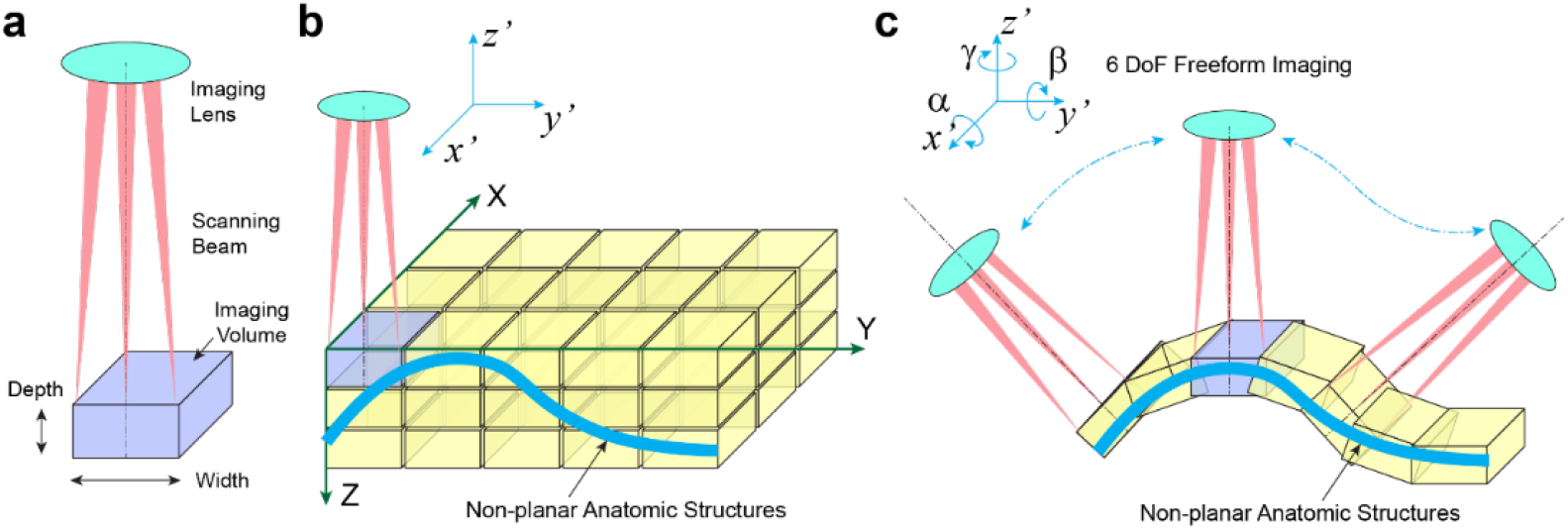
Freeform imaging for expanding the optical FOV. a) The optical FOV for conventional microscopy is determined by the DOF. b) Mechanical translation along the XYZ axes can be used to expand the optical FOV. The blue box shows the optical FOV for a single volumetric acquisition, and the yellow boxes show the space in which the FOV can be expanded. c) Freeform robotic scanning increases the FOV and changes the orientation of the sample arm relative to the sample.

We developed a freeform robotic OCT system to meet these needs. This system integrates an OCT sample arm with a six-axis robotic arm, allowing fully flexible, conformal scanning across non-planar sample geometries (Fig. 1c). The inclusion of rotational motion enhances the system’s adaptability to maintain a normal OCT illumination to an arbitrary sample surface, thereby optimizing imaging quality. Additionally, tailored conformal CIV to non-planar geometries effectively reduces image acquisition and processing time.

Using a six-axis robotic arm offers flexibility in acquiring conformal CIV but introduces two primary technical challenges. First, the conformal CIV is specific to sample geometry, requiring knowledge of the sample geometries’ local coordinates relative to the robotic arm’s reference frames. Second, despite the high precision of the six-axis robotic arm, its accuracy is relatively limited ^10-14^, especially compared with the resolution of OCT. This discrepancy poses significant challenges in reconstructing CIVs by montaging multiple OCT volumetric images with an axial resolution of 2-5 μm. While the top commercial robots can achieve positioning precision of 5 μm, their accuracy falls behind at approximately 200 μm. Bridging this gap of nearly two orders of magnitude between spatial accuracy and the spatial resolution of OCT presents a grand challenge in reconstructing CIVs. Even though the positioning accuracy of 6-axis robots can be improved with high-precision metrology tools like coordinate measurement machines and laser interferometer trackers^15,16^, these approaches add unwanted burdens regarding technical complexity and cost, particularly in imaging biological samples.

To address these challenges, we repurposed the OCT as a built-in optical metrology tool, achieving two critical objectives simultaneously. Firstly, we delineated the distinct geometries of the samples, aligning the robotic arm’s reference frames with the local coordinates of the samples. Secondly, we identified unique anatomical features on the samples using OCT, guiding the precise reconstruction of CIVs through the montage of multiple OCT volumetric images. This process was further augmented by a new algorithm based on simultaneous localization and mapping (SLAM), tailored for freeform OCT. We tested our freeform OCT by imaging the anterior segment and aqueous humor outflow (AHO) pathway in mouse eyes as a model system.

We selected the AHO pathway for its anatomical complexities, making imaging the entire AHO pathway with traditional OCT difficult, and its clinical relevance to glaucoma. The main risk factor for glaucoma, a leading cause of irreversible blindness globally, is elevated intraocular pressure (IOP).^17^ The equilibrium between aqueous humor inflow and outflow through the AHO pathway is crucial for IOP regulation. In both rodents and humans, the trabecular AHO pathway, comprising the anterior chamber, trabecular meshwork (TM), Schlemm’s canal (SC), collector channels (CC), and distal outflow vessels, is responsible for the majority of aqueous drainage.^18^ Pathological increases in resistance within the trabecular AHO pathways lead to elevated IOP. Visualizing the AHO pathway’s anatomy in three dimensions (3D) at a microscopic spatial resolution can revolutionize glaucoma drug development^19,20^ and enhance the efficacy of minimally invasive glaucoma surgeries (MIGS) in patients^21,22^. However, imaging the AHO pathway presents substantial challenges using existing OCTs due to the AHO pathway’s anatomical complexity, including fine features with dimensions as minute as 2 μm, the orientation of those features oblique to the optical axis of the eye, and its circumferential distribution around the limbus. Additionally, CCs and distal pathways radiate outward, extending several millimeters beyond the limbus. These distinctive anatomical characteristics underscore the necessity for a freeform robotic OCT system.

We developed a robotic visible-light optical coherence tomography (vis-OCT) to image the entire AHO and its key components (TM, SC, and CC) within the CIV. Vis-OCT achieved an axial resolution of 1.3 μm, well suited for visualizing AHO pathways in mice. During the circumferential robotic scanning, we ensured that the vis-OCT illumination light remained conformal and perpendicular to the limbal surface. Our OCT-based SLAM achieved an accuracy below 10 μm, overcoming the intrinsic limit of the robotic arm’s limited accuracy. This enabled the volumetric reconstruction of the entire AHO pathway, which further unveiled segmental variations in SC morphology and CC distribution and offered new insights into the structural complexity of the AHO system.

## Results

### Robotic vis-OCT system and data acquisition workflow

Fig. 2a shows the schematic of the robotic vis-OCT system. The filtered output from a supercontinuum laser was coupled into a 90:10 fiber coupler (FC1), with 10% output to the sample arm and 90% output to the reference arm. In the sample arm, collimated light from the fiber coupler was scanned by a galvanometer and focused onto the sample. Back-scattered light from the sample passed through FC1 and interfered with the light from the reference arm in the 50:50 fiber coupler (FC2). Two well-calibrated spectrometers detected the interferogram simultaneously for balanced detection^23^. We designed a compact vis-OCT sample arm (white dashed square, Fig. S1a) and mounted it on a robotic arm (Meca 500, Mecadamic Inc.), which supports a payload of 0.5 kg with a positioning precision of 5 μm over a maximum reach of 225 mm. The sample arm enclosure was 3D printed using carbon fiber-reinforced polycarbonate polymer materials to reduce the total weight to 0.29 kg. We integrated a pupil camera (outer diameter: 2 mm) and a time-of-flight (TOF) proximity sensor to the sample arm. The pupil camera provided the initial video guidance to locate the eye, and the TOF sensor prevented the sample arm from colliding with the eye.

**Fig. 2.**
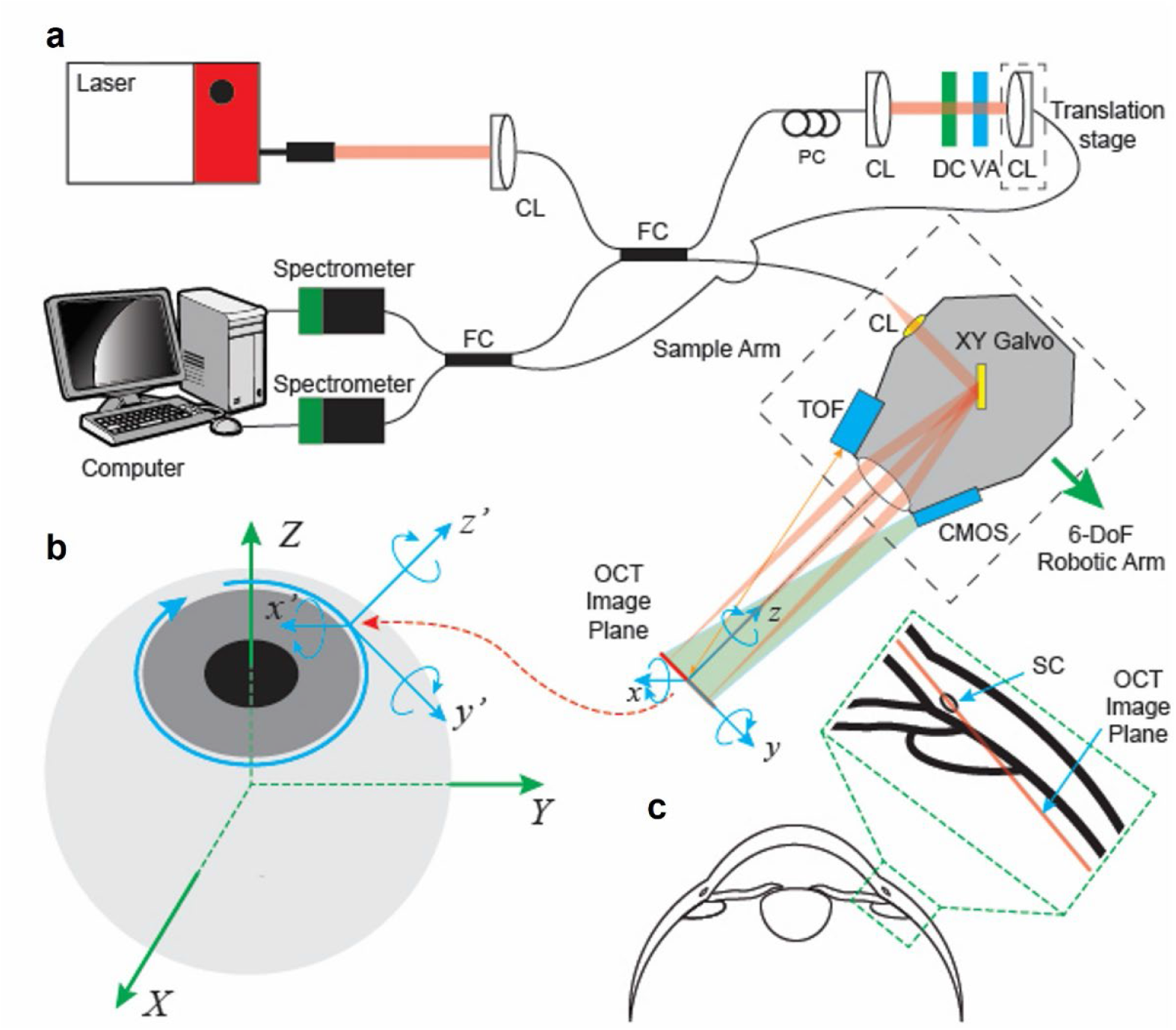
Experimental setup of robotic vis-OCT and imaging geometry. a) Schematic of vis-OCT system. CL: collimator; FC: fiber coupler; PC: polarization controller; DC: dispersion compensation; VA: variable attenuator; TOF: time-of-flight sensor; CMOS: pupil camera. The sample arm, inside the dotted black box, is attached to a 6-DoF robotic arm. The imaging coordinates of the OCT is the tool reference frame (TRF), with the z-axis parallel to the incident OCT beam and the OCT image plane coinciding with the xy plane of the TRF. b) Imaging geometry around the limbus of the eye and the relationship between two reference frames: the eye reference frame (ERF) represented by the XYZ-coordinate (green) and the TRF of the robot represented by the xyz-coordinate (blue). Following calibration, 360-degree freeform imaging of the SC is achieved by rotating the TRF around the Z-axis of the ERF. During this rotation, the origin of the TRF follows a blue-colored circular path while maintaining the z-axis of the TRF perpendicular to the eye surface. This alignment ensures that the OCT image plane remains tangent to the surface of the eye. c) Cross-sectional view of the anterior segment, with a magnified view of the SC shown in the dashed green box.

Operating the robotic vis-OCT system requires managing two independent cartesian coordinate systems of the eyeball and the vis-OCT imaging volume (Fig. 2b.) The eyeball coordinates are represented by the eye reference frame (ERF, X-Y-Z axes), where the Z-axis aligns with the eyeball’s visual axis, and the origin is positioned at the center of the spherical surface approximating the cornea. The vis-OCT imaging volume’s coordinates are represented by the tool reference frame (TRF, x-y-z axes). Here, the x-y plane coincides with the vis-OCT image plane. The orientation of the x-axis and y-axis is calibrated to align with the scanning directions of the galvanometer scanner. The origin of TRF is at the center of the vis-OCT FOV, with the z-axis pointing along the optical axis of the sample arm. For conformal imaging of the AHO pathways, it is important to accurately calibrate the 6-DoF coordinate transformation between the ERF and the TRF. A detailed calibration process is provided in the Supplemental Methods 2 and 3.

To perform conformal AHO pathway imaging after calibrating the coordinate transformation from TRF to ERF, we first rotated the TRF 60 degrees about either the ERF X or Y-axes, which is near the angle of the trabecular AHO pathway relative to the ERF. Then we adjusted the OCT sample arm to rotate the TRF 360 degrees around the ERF’s Z-axis (path highlighted by the red-colored trajectory in Fig. 2b). For each sample arm position, the OCT illumination light along the z-axis of the TRF remained normal to the tissue surface (Fig. 2c).

### Influence of vis-OCT illumination incident angle

A key advantage of robotic OCT is its capability to adjust incident angle as needed. We examined the influence of illumination incident angle in imaging AHO pathway based on the contrast-to-noise ratio (CNR). Since the axial resolution of nearly all OCT is several folds better than its lateral resolution, anatomical features are best resolved axially instead of laterally^24^. Hence, maintaining a normal illumination can maximize the benefit of vis-OCT’s high axial resolution in imaging fine anatomical features in the AHO pathway, such as the SC. If we simplify the cross-section of the SC as an ellipse, where the minor axis corresponds to its smaller dimension and the major axis to its larger dimension, a normal OCT illumination to the limbus will align with the minor axis to best resolve SC.

We imaged the SC (Fig. 3a) and the cornea (Fig. 3b) using two vis-OCT illumination incident angles: normal to the major axis of SC (highlighted by the red arrows) and to the central cornea (highlighted by the blue arrow). When the vis-OCT illumination aligns with the normal axis of the cornea, it was difficult to resolve the borders of SC (blue box in Fig. 3c) and hard to visualize the limbal vasculature from the vis-OCT angiography or vis-OCTA (Fig. 3d). We differentiated the corneal layers in the central cornea (blue box in Fig. 3e) but not the peripheral cornea (red box in Fig. 3e). When the vis-OCT illumination aligns with the minor axis of SC, we well resolved the borders of SC (red box in Fig. 3f) and the limbal vasculature network from vis-OCTA (Fig. 3g). Additionally, we observed different corneal layers in the peripheral cornea (red box in Fig. 3h) but not the central cornea (blue box in Fig. 3h).

**Fig. 3.**
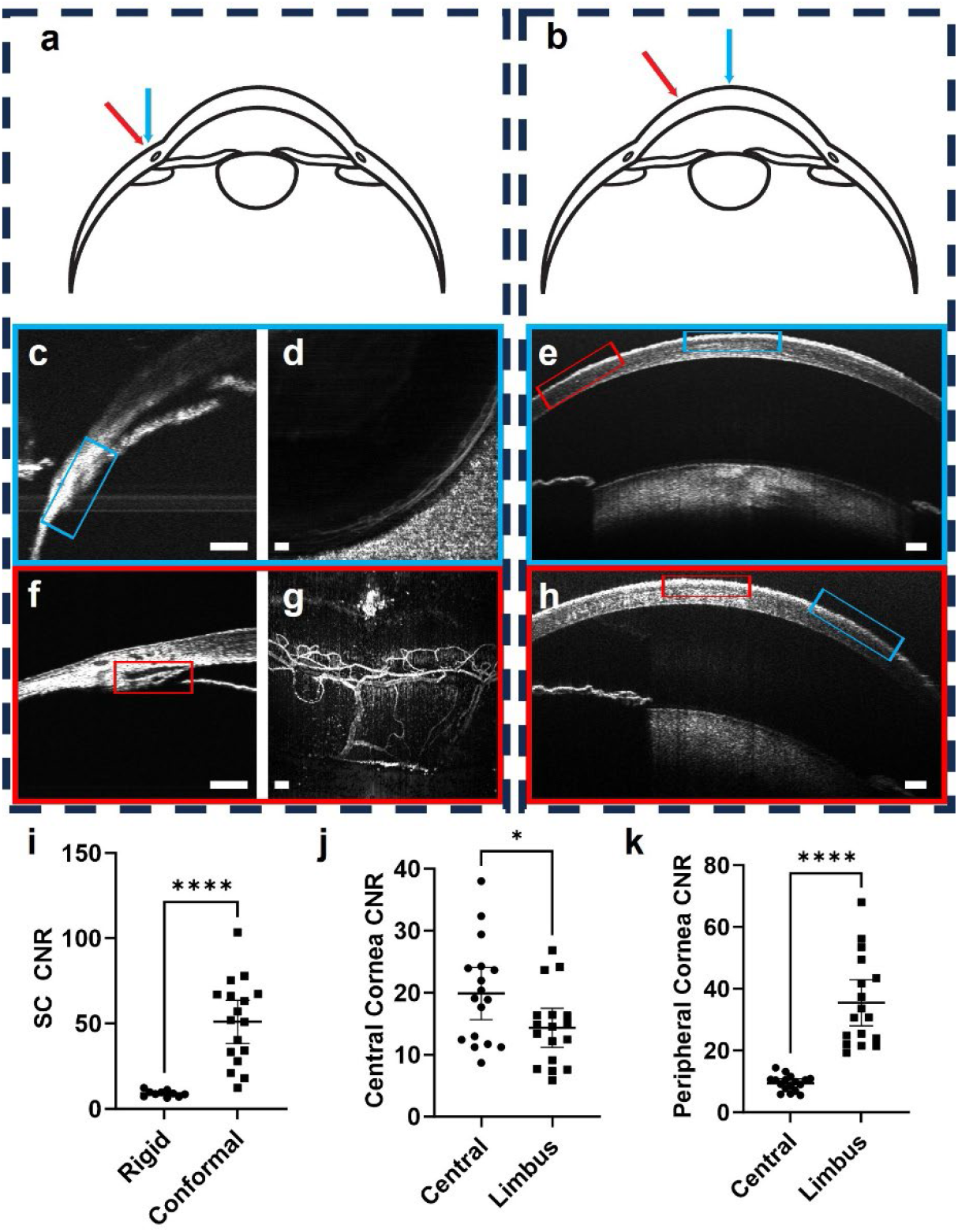
Effects of vis-OCT illumination angle. a) Imaging the limbus region at two distinct illumination angles: normal to the central cornea (blue arrow) and normal to the limbus (red arrow). b) Imaging the central cornea at two distinct illumination angles: normal to the central cornea (blue arrow) and normal to the limbus (red arrow). c) B-scan image of SC (blue box) using the illumination angle highlighted by the blue arrow in panel a. d) Vis-OCTA of the limbal vasculature using the illumination angle highlighted by the blue arrow in panel a. e) B-scan image of the cornea using the illumination angle highlighted by the blue arrow in panel b. f) B-scan image of SC (red box) using the illumination angle highlighted by the red arrow in panel a. g) Vis-OCTA of the limbal vasculature using the illumination angle highlighted by the red arrow in panel a. h) B-scan image of the cornea using the illumination angle highlighted by the red arrow in panel b. i) The CNR between SC and surrounding tissue is greater with beam orientated normal to the limbus (conformal) than the central cornea (rigid). j) CNR between the epithelium and stroma in the central cornea is greater when the illumination beam is normal to the central cornea. k) CNR between the epithelium and stroma in the peripheral cornea is greater when the illumination beam is normal to the limbus. All scale bars are 100 μm.

Quantitatively, the CNRs between SC and surrounding tissue were 9.0 ± 1.8 when the vis-OCT beam was normal to the central cornea and 51.0 ± 24.7 when orientated along the minor axis of SC (Fig. 3i). The CNRs between the epithelium and stroma in the central cornea were 19.9 ± 8.2 when orientated normal to the central cornea and 14.3 ± 6.1 when orientated along the minor axis of SC (Fig. 3j). The CNRs between the epithelium and stroma in the peripheral cornea were 9.5 ± 2.5 when orientated normal to the central cornea and 35.5 ± 14.4 when orientated along the minor axis of SC (Fig. 3k). In conclusion, best imaging quality requires vis-OCT illumination beam to be normal to the sample surface, requiring conformal scanning.

### SLAM-based volumetric registration for robotic vis-OCT

To reconstruct a CIV of the anterior segment of mouse eyes, we acquired eight vis-OCT volumes separated by 45 degrees angular increments along the trajectory illustrated in Fig. 2b. For each volume, we recorded the orientation of the TRF relative to the ERF and reconstructed both the microanatomy and angiogram (Fig. 4a). After we determined which volumes overlapped with each other based on the recorded TRF orientations, we registered and montaged adjacent volumes to reconstruct the entire CIV. To montage adjacent volumes, we segmented the tissue surface and represented the outer surface as a point cloud ^25^ (step 1 in Fig. 4b). We identified common vessel branch points between adjacent volumes, extracting the outer surface at the lateral position of the branch points (step 2 in Fig. 4b). Then, we used the M-estimator Sample Consensus algorithm^26^ to determine the rigid geometric transformation between the common branch points (step 3 in Fig. 4b). After determining the rigid geometric transformation between overlapping regions on adjacent point clouds, the mean distance between each point on a point cloud and the closest point on the adjacent point cloud was 9.6 ± 2.3 μm. All point clouds were transformed into the ERF coordinate system. Following transformation into the ERF, we found the overlapping regions of the point clouds (Fig. 4c) and used the iterative closest point algorithm^27^ to refine the transformations of each reference frame relative to the ERF. More details of the SLAM are provided in Supplemental Methods 4.

**Fig. 4.**
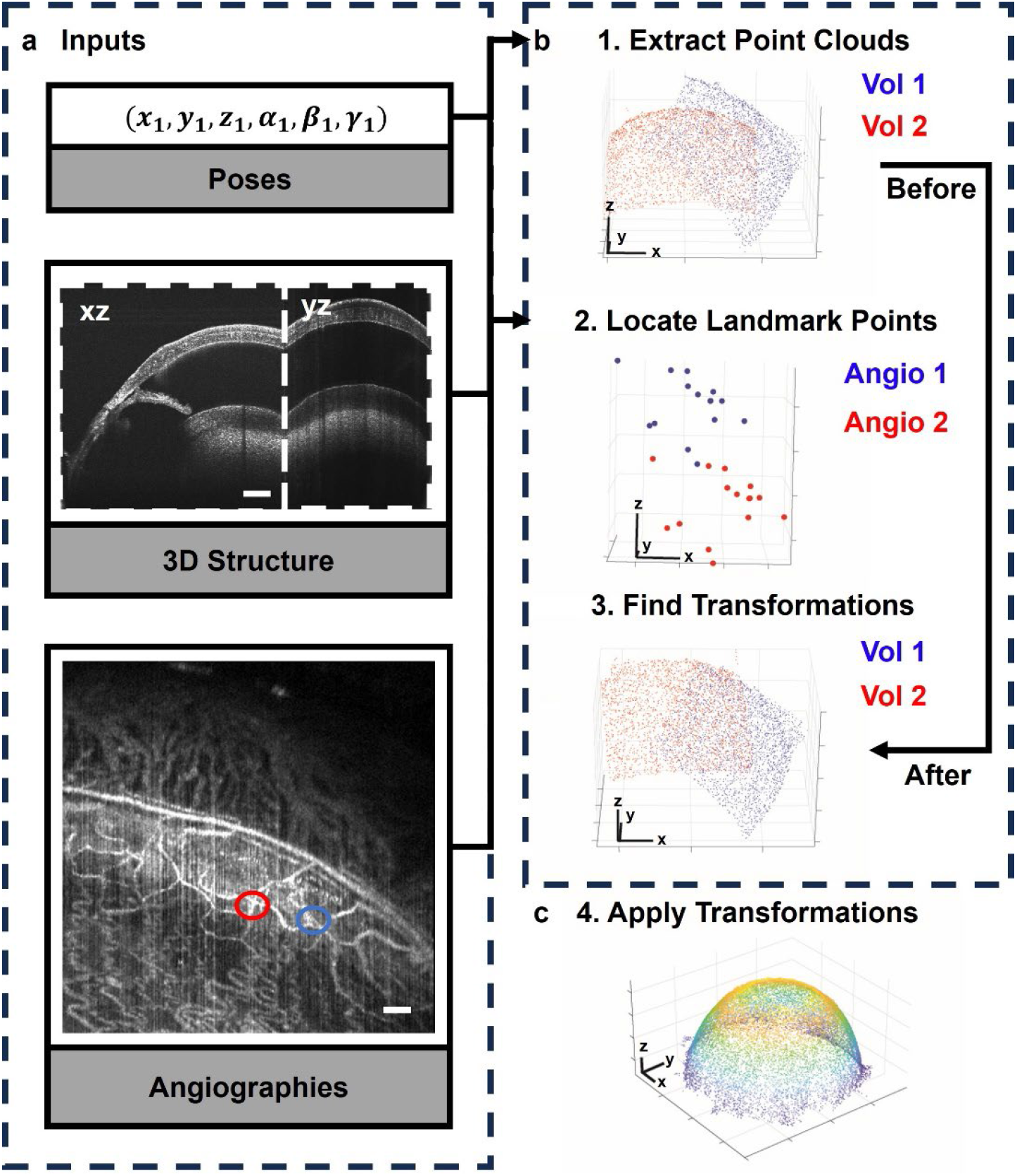
Workflow of montaging multiple vis-OCT volumes into a CIV. a) Input data acquired by freeform robotic OCT. Poses estimate the acquired volume’s location and determine which volumes overlap. Each acquired volume consists of a 3D structure and angiographic data. b) Landmark points from overlapping volumes are used to merge the volumes. (1) The outer surface of the volumetric data is represented as a point cloud; (2) landmark points corresponding to vessel branch points are extracted from the point clouds. Two examples of landmark points are given by the blue and red circles in (a); (3) Common landmark points between volumes are registered to each other to determine the transformation between overlapping volumes. c) Following the calculation of the relative transformations between volumes, all volumes are mapped into the ERF. A modified iterative closest point algorithm addressed the loop closure problem. Scale bars for volume data are 200 μm, and for angiographic data are 100 μm.

### Reconstructing the anterior segment

Following SLAM, we applied the calculated geometric transformations to each vis-OCT volume and visualized the reconstructed anterior segment from the anterior (Fig. 5a) and posterior (Fig. 5b) perspectives. A sample cross-section located 560 μm from the center of the eye shows a smooth corneal and iris surface, indicating accurate montaging (Fig. 5c). We transformed the coordinate system of the reconstructed volume from a cartesian coordinate into a spherical coordinate system and identified the mean structural and angiographic signal as a function of *θ* and *ϕ*. Following coordinate transformation, we projected the structural and angiographic volumetric data into two dimensions using a stereographic projection (Figs. 5d-5e)^28^. Features, including the trajectory of individual blood vessels along the projections, are smooth and continuous between the montaged volumes.

**Fig. 5.**
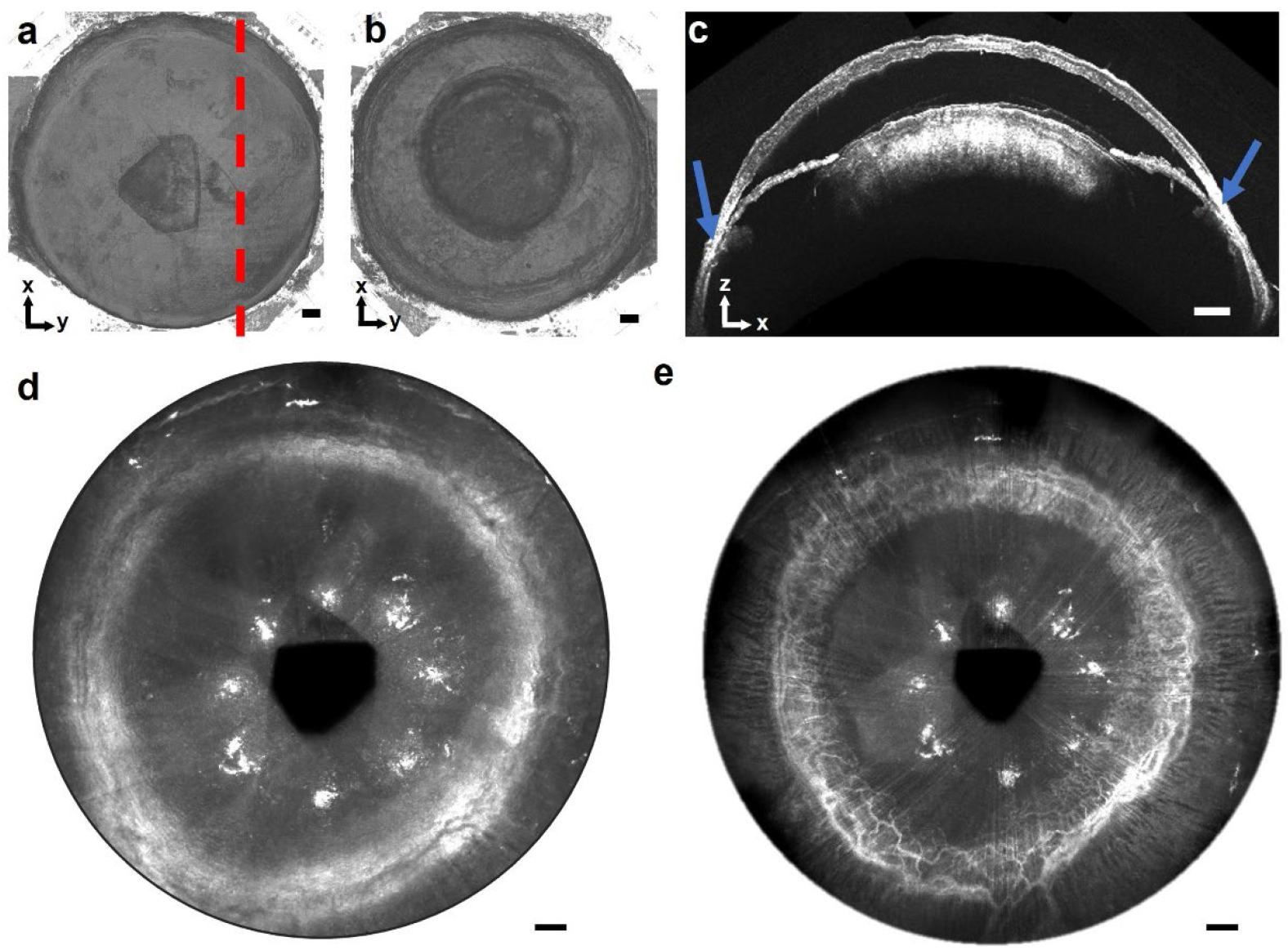
Reconstructed CIV of the entire anterior segment. a) Volumetric reconstruction of the anterior segment viewed from the anterior direction, visualizing the outer surfaces of the cornea and limbus. b) Volumetric reconstruction of the anterior segment viewed from the posterior direction, depicting the lens and inner surface of the limbus. c) A representative cross-sectional image along the red line in panel a, with the blue arrows indicating the SC. d) Steographic projection of structural vis-OCT volumetric data. e) Stereographic projection of the vis-OCTA volumetric data highlighting connected and continuous limbal vasculature. Scale bars are 200 μm.

### Segmental pattern of SC morphology

From the reconstructed anterior segment, we segmented the SC around the entire globe (Fig. 6a), which forms a continuous lumen (Fig. 6b). We approximated the cross-section of SC by an ellipsoid, with the major and minor axes corresponding to the SC width and height, respectively and selected several planes to visualize SC around the 360 degrees of the eye. When visualized along a plane passing through major axes of each cross-section of SC, SC forms a continuous hypo-reflective ring immediately lateral to the anterior chamber (Fig. 6c). Furthermore, we visualized segmental SC height patterns after digitally resampling^29^ (see Supplemental Methods 5) the anterior segment along the lateral center of SC (Fig. 6d). Overall, SC height is larger in the nasal and temporal quadrant for a C57BL/6 mouse. For each mouse, we measured segmental patterns in SC area, height, and width as a function of angle around the globe (Figs. 6e-6g).

**Fig. 6.**
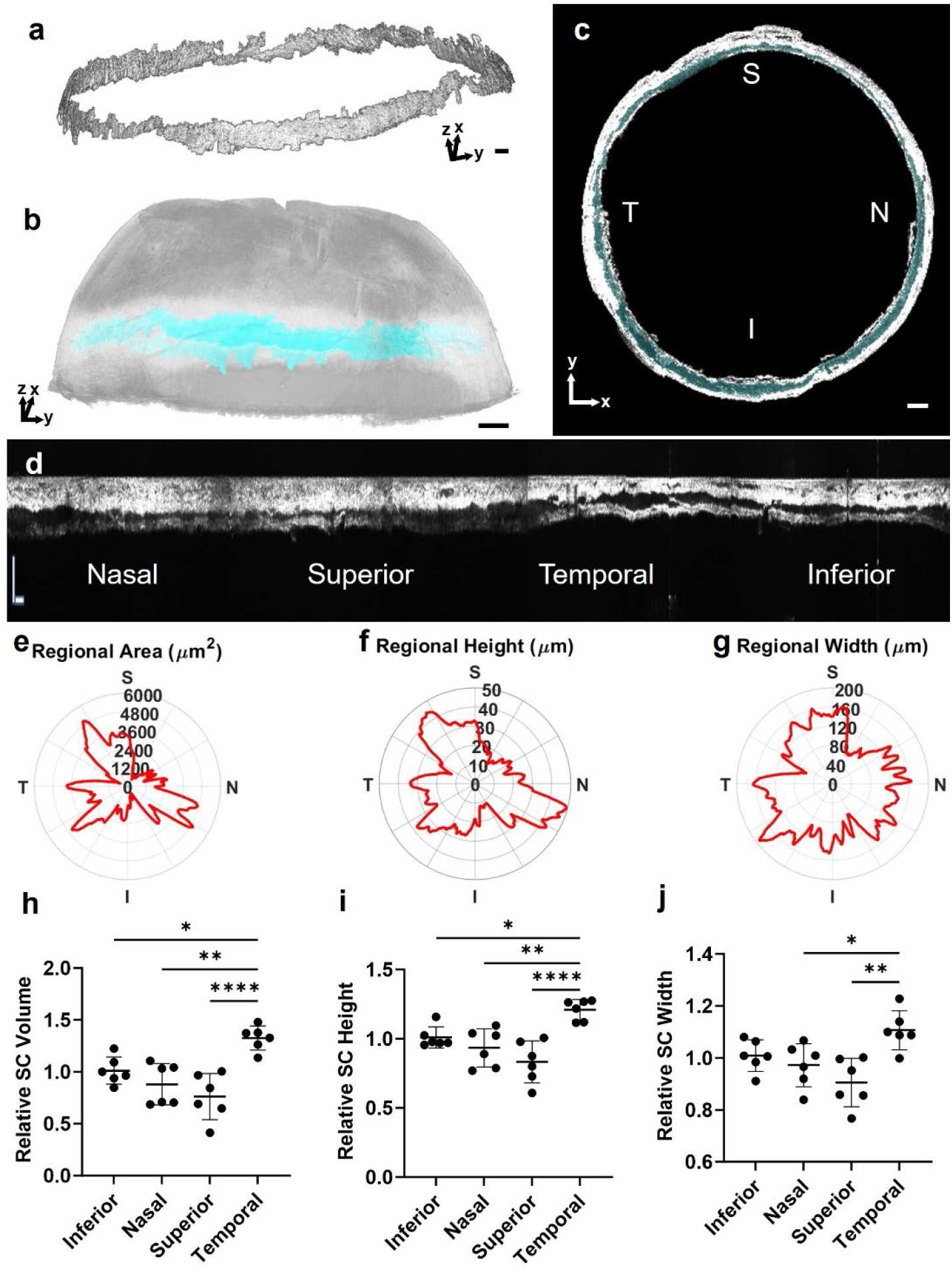
Circumferentially segmental patterns of SC morphology. a) A 3D reconstruction of SC.b) SC pseudocolored in light blue within the structural 3D reconstruction of the anterior segment.c) Projection of SC along its major axis circumferentially. The position of SC is overlayed in cyan.d) An unwrapped view of SC along its circumference divided by the four quadrants. e) Polar distribution of SC cross-sectional area as a function of circumferential orientation. f) Polar distribution of SC height as a function of circumferential orientation. and g) Polar distribution of SC width as a function of circumferential orientation. h) Relative SC volume for each quadrant compared to the average across all quadrants. i) Relative SC height for each quadrant compared to the average across all quadrants. j) Relative SC width for each quadrant compared to the average across all quadrants (n = 6). * indicates p < 0.05, ** p < 0.01, **** p < 0.0001. Scale bars in (a)-(c) are 200 μm and (d) is 100 μm.

Relative to the mean volume across all quadrants for each eye, SC volume was 32.6 ± 11.7% larger in the temporal quadrant, 23.8 ± 22.2% smaller in the superior quadrant, 12.0 ± 19.9% smaller in the nasal, and 1.2 ± 13.1% larger in the inferior quadrant. Volume in the temporal quadrant was greater than the other quadrants (Fig. 6h). Relative to the mean height across all quadrants, SC height was 21.0 ± 7.4% larger in the temporal quadrant, 16.7 ± 15.2% smaller in the superior quadrant, 6.6 ± 13.8% smaller in the nasal, and 0.9 ± 7.7% larger in the inferior quadrant. Height in the temporal quadrant was greater than the other quadrants (Fig. 6i). Relative to the mean width across all quadrants, SC width was 10.7 ± 7.5% larger in the temporal quadrant, 9.5 ± 9.3% smaller in the superior quadrant, 2.8 ± 8.3% smaller in the nasal, and 0.9 ± 6.1% larger in the inferior quadrant. Width in the temporal quadrant was greater than the nasal and superior quadrants (Fig. 6j).

### Segmental distribution of CCs

As the segmental distribution of CCs has been hypothesized to influence MIGS outcomes^30^, we visualized each CC around the globe in mice from the reconstructed anterior segment. We mapped the positions of CCs around the eye and correlated locations with the local SC area in C57BL/6 mice (Fig. 7a). We found that the mean total number of CCs was 33.7 ± 6.5. The average number of CC in the inferior, nasal, superior, and temporal quadrants are 6.2 ± 2.0, 10.5 ± 2.8, 6.3 ± 2.3, and 10.7 ± 2.6 respectively (Fig. 7b). Overall, the nasal and temporal quadrants contained more CCs than the inferior and superior quadrants (p <0.05).

**Fig. 7:**
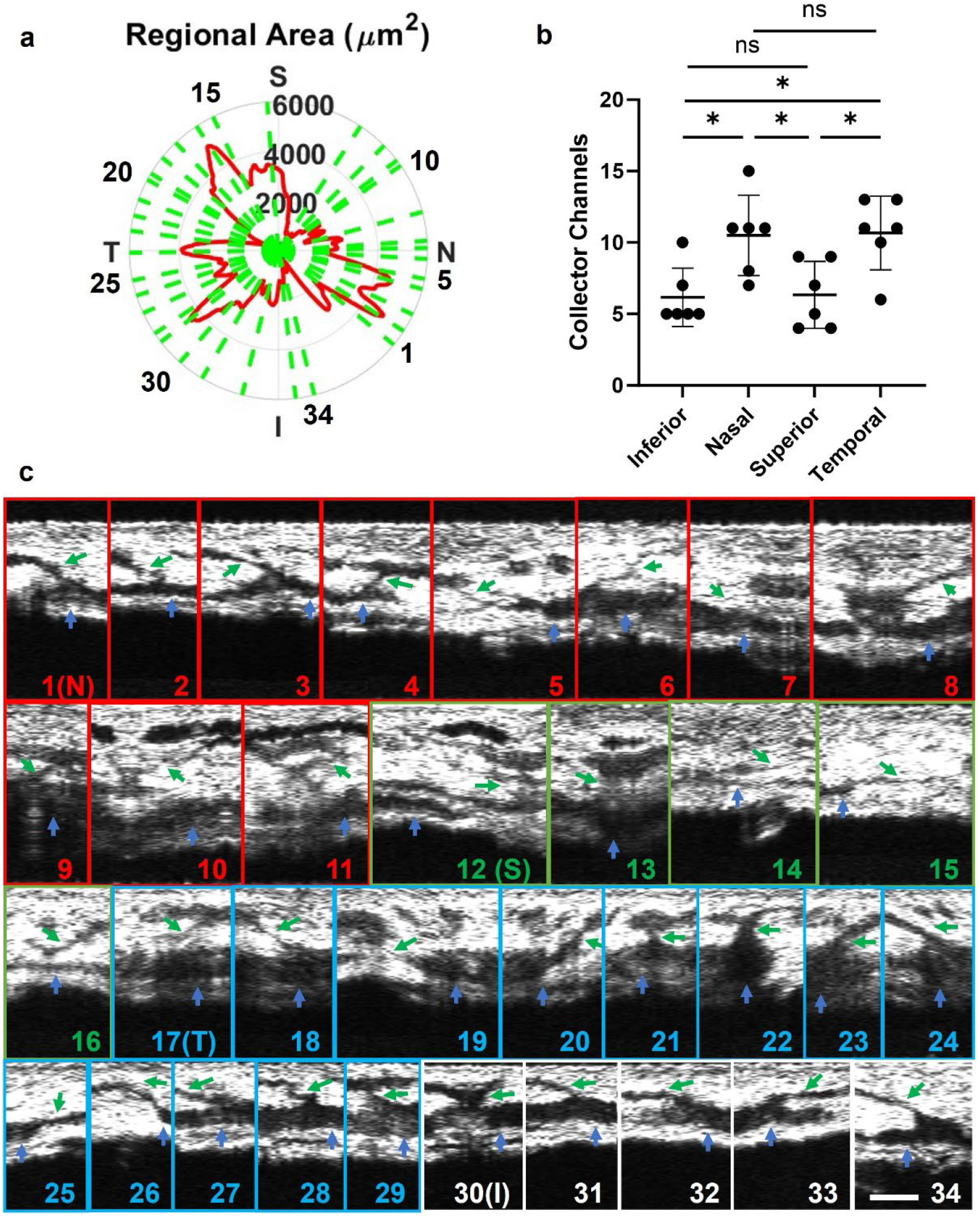
Circumferentially segmental patterns of CCs. a) Polar distribution of SC area and CC positions as a function of circumferential orientation in a typical C57BL/6 mouse. The red curve represents the SC area, and the green dotted lines indicate the CC position. b) Distribution of CC per quadrant. c) Magnified view of all CCs in panel a. The blue arrow points to SC, and the green arrows to CC. The numbering of CC corresponds to the numbering of green lines in panel a. CCs labeled in red are in the nasal quadrant, green in the superior quadrant, blue in the temporal quadrant, and white in the inferior quadrant. The scale bar is 50 μm.

After digital resampling, we visualized every single CC in a C56BL/6 mouse (Fig. 7c). For each resampled CC image, we highlighted the connection between SC (blue arrows) and CC (green arrows). Several different CC morphologies were observed, with some CCs open throughout (#30 in Fig. 7c), some CCs exhibiting pinch points (#6 in Fig. 7c), some CCs exiting at a vertical angle (#22 in Fig. 7c), and some CCs exiting at a side angle (#33 in Fig. 7c). These results further validated the complexity of AHO pathway and the need for conformal OCT imaging.

## Discussion

Limitations in mechanical scan angle and illumination orientation constrain the imaging volume for most OCT systems. Here, we present a robotic OCT capable of freeform eye imaging to overcome these limitations. Robotic OCT has previously been used to image the retina but has not been utilized to expand the optical FOV or image the AHO pathway^5,31-33^. We developed a volume registration algorithm based on 3D SLAM for OCT, generating a CIV that encompasses the entirety of the AHO pathway in mice.

Compared with stationary OCTs, our robotic OCT has complete flexibility in controlling illumination incident angles during data acquisition. We found that CNR in imaging the AHO and cornea depended on OCT’s illumination incident angle. Optimal imaging of complex non-planar tissues occurs when the OCT illumination beam is normal to the tissue surface. In our case, we accomplished this by calibrating the ERF relative to the TRF and rotating about the z-axis of the ERF (see Supplemental Methods 2-3). Using conformal scanning, we resolved the SC with much higher CNRs than could be achieved with a fixed beam angle. One advantage of conformal scanning is that the sample does not need to be repositioned to achieve normal illumination. For *in vivo* imaging, this will allow fewer motion artifacts and improved reproducibility^34-36^.

Following acquisitions of OCT volumes from multiple poses around the eye, we represented the surface of each volume as a point cloud and performed SLAM to integrate all the volumes. In traditional robotics, SLAM integrates robotic positional data with images of the environment to map out unknown environments^37^. In most cases, robotic positional sensors are more accurate than the surrounding objects being reconstructed. In our case, the resolution of OCT is several folds higher than the accuracy of robotic arms. Our vis-OCT has an axial resolution of 1.3 μm while the accuracy of most precise robotic arms exceeds 100 μm^38^. Consequently, robotic positional sensors cannot reliably merge all vis-OCT volumes. To address this issue, we developed a montaging method integrating robotic positional data with acquired OCT volumes to reconstruct the anterior segment of the eye. Robotic poses were only used to determine whether two volumes have sufficient overlapping regions. The fine montaging of 3-D structural features was accomplished by optimizing the geometric transformations mapping the OCT landmark points of adjacent point clouds onto each other with under 10 μm distance between landmark points in adjacent volumes, an orders of magnitude better than the accuracy of the robotic arm. The outer surface of the merged volumes and 360-degree reconstructed SC was smooth and continuous, indicating the accuracy of volumetric montaging.

The integration of mechanical design and novel processing expands the CIV of OCT, enabling the reconstruction of complex non-planar anatomies. Nearly all existing OCT montaging algorithms register en-face projection images^39,40^. However, en-face projections become nonlinearly distorted relative to each other when their curvature relative to the incident beam is significantly different from each other, as in our situation. To circumvent this obstacle, we used 3D volumetric surface point cloud data to perform SLAM for OCT, which does not suffer the same angular distortion.

Finally, we demonstrate the utility of an increased CIV by imaging the full 360 degrees of the AHO pathway, which may improve clinical glaucoma management. From a scientific perspective, reconstructing AHO pathways can improve our understanding of how outflow resistance is regulated. One prevalent question is the root cause of resistance distal to SC^41^, which may be caused by pinch points in the distal pathway. We observe collapsed regions of the distal pathway in our resampled CC, which is consistent with the existence of pinch points. From a surgical perspective, it has been demonstrated that segmental patterns in anatomy, outflow, and CC distribution influence the success of MIGS^21,30,42-44^. We quantified segmental differences in SC size and CC distribution and found SC was the largest in the temporal quadrant, and the temporal and nasal quadrants had the most CCs. Although more investigation is necessary on guiding surgical decisions, this work lays the technological groundwork for this goal.

## Materials and Methods

### Robotic vis-OCT system

Broadband light is emitted from a supercontinuum laser (SuperK 78 MHz EXW-6, NKT Photonics). The output laser light passed through a dichroic mirror (DMSP650, Thorlabs), bandpass filter (FF01-560/94-25, Semrock), and a spectral shaping filter (Hoya B-460, Edmund Optics). The incident light was coupled into a 90:10 fiber coupler (TW560R2A2, Thorlabs).

The sample arm of our vis-OCT was attached to a high-precision six-axis robotic arm (Meca500, Mecademic Inc.), which has a repeatability of 5 μm^45^. The sample arm consisted of a collimator (RC02APC-P01. Thorlabs), a 3-mm compact galvanometer (RC025547, ScannerMAX), and a 25-mm achromatic lens (Ac127-025-A, Thorlabs). The reference arm consists of 2 collimators (HPUC-2A3A-400/700-P-6AC-11, OZ Optics), BK7 dispersion glass (WG11010, Thorlabs), a polarization controller (FPC030, Thorlabs), and a variable attenuator (NDC-50C-4M, Thorlabs). Light from the sample and reference arm are coupled into a 50:50 fiber coupler (TW560R5A2, Thorlabs). After the couplers, the interference pattern was sent to 2 spectrometers (Blizzard SR, Opticent Inc.).

### Image acquisition protocol

Additional details regarding the data acquisition are found in Supplemental Methods 2&3.

#### Defining the ERF

We brought the eye into the OCT field of view, verifying the eye position with the pupil camera (Supplemental Methods 3). We developed a customized LabVIEW program (2021) to preview real-time B-scans and perform image analysis of B-scans simultaneously. We applied customized algorithms to align the z-axis and transverse origin of the ERF and TRF (Supplemental Methods 3). We completed this process for both the x and y-axes of the TRF separately.

#### 360-degree scanning

After defining the ERF, we rotated the TRF 60 degrees about the WRF X-axis. Afterward, eight OCT volumes were sequentially acquired after 45-degree rotations of the TRF around the WRF Z-axis

The spectral range of our vis-OCT system was 510 nm – 610 nm. For each eye, we acquired eight vis-OCT volumes orientated 45 degrees from each other. The incident power on the eye was 1 mW, and the A-line rate was 75kHz. The vis-OCT’s lateral FOV was 2.04 mm. We applied a temporal speckle averaging^46^ data acquisition pattern to acquire each volume, where each B-scan was acquired twice, and three volumes were acquired. B-scans from each volume were subsequently averaged.

### Data processing and measuring SC parameters

We reconstructed the anatomical vis-OCT images following methods in our previous work^47^. Further, we processed the angiography data using published OMAG and SSADA algorithms^48,49^. After reconstructing each volume, we mapped all volumes into the ERF using our SLAM algorithm. Additionally, we segmented the entire SC around the globe. For segmentation, we selected the lumen corresponding to SC in each B-scan of every volume and applied a flood fill algorithm to identify the borders of the lumen. We manually checked and applied corrections to all segmented B-scans using MATLAB’s (2022) Volume Segmenter package. By summing all SC cross-sectional areas within each quadrant of the eye, we calculated the regional volume per quadrant of the eye. Once SC was segmented, SC height and width corresponded to the minor and major axis dimensions of an ellipse fit to SC^50^. See Supplemental Methods 4 and 5 for more information regarding volumetric montaging and outflow pathway reconstruction.

### Animal care and handling

We anesthetized all mice by intraperitoneal injection (10 mL/kg body weight) of ketamine/xylazine cocktail (ketamine 11.45mg/L, xylazine 1.7mg/L in saline). While under anesthesia, the mouse’s body temperature was maintained using an infrared lamp. To expose the entire 360 degrees of the limbus, we made a small incision to the temporal and nasal eyelids and inserted a speculum underneath the eyelids (see Supplemental Methods 6). All experimental procedures were approved by the Institutional Animal Care and Use Committee (IACUC) at Northwestern University.

### Statistical analysis

All statistical comparisons, unless otherwise specified, were done after performing a one-way ANOVA using GraphPadPrism 9.5.1. When multiple comparisons were made, we performed a Tukey multiple comparison correction. We used the F-test to determine whether the fit slopes of any fit line were significantly different from zero.

## Supporting information

Supplemental Methods and Results

## Acknowledgments

This work was supported in parts by National Institutes of Health grants R01EY029121, U01EY033001, R01EY033813, R01EY034740, R01EY034353, R01EY030501, F30EY034033, and R44EY026466, Illinois Society for the Prevention of Blindness, and the Christina Enroth-Cugell and David Cugell Fellowship.

