## Supplemental Methods and Results for "Freeform robotic optical coherence tomography beyond the optical field-of-view limit"

### Supplemental Information

#### Supplemental Discussion 1: Robotic Vis-OCT and Definition of Reference Frames

We enclosed the sample arm of our vis-OCT in a compact black 3D-printed chassis (white dashed square, Fig. S1a), which we attached to a six-axis Meca500 robotic arm. Our sample arm weighs ~0.29 kg, less than the specified Meca500 robotic arm payload of 0.5 kg[1]. The 6 degrees of freedom (DoF) enables the robotic arm to achieve any translational (x,y,z) and angular (α, β, γ) orientation within the robotic arm's range of motion.

Every object is associated with a reference frame, which consists of a 6DoF Cartesian coordinate system. Objects are capable of translating and rotating about the 3 axes of a 6DoF reference frame. The pose, which is the position and orientation of an object and its reference frame relative to another reference frame, is given by (x, y, z, α, β, γ). Under this convention, x, y, and z define the relative positions between the origins of the reference frames. Similarly, α, β, γ refer to the relative orientation between the reference frames and follow the XYZ Euler angle convention. When describing the pose of a reference frame $F_1$ relative to $F_0$, a frame initially aligned with $F_0$ rotated by α degrees around the x-axis of $F_0$, then β degrees about the y-axis, and γ degrees about the z-axis will align its axes with $F_1$.

Our robotic arm has several reference frames (Fig. S1b), whose poses relative to each other are defined. The eye reference frame (ERF) and tool reference frame (TRF) are the most important for our purposes. The ERF is the coordinate system of the eye being imaged, and the TRF is the coordinate system of the vis-OCT imaging volume. Poses for the ERF are given by XYZRPW. For the ERF, the Z-axis is the visual axis of the eye, and RPW corresponds to rotations about XYZ. The X and Y-axes of the ERF extend in the nasal/temporal or superior/inferior directions depending on the pose of the eye to the initial OCT imaging volume. The origin of the ERF is the centroid of a sphere approximating the eye's corneal surface. The poses for the TRF are given by xyzrpw. For the TRF, the xy axes correspond to the scanning directions of the galvanometer, and the z-axis corresponds to the primary optical axis. The origin of the TRF, known as the tool center point (TCP), is the focal point of the OCT scanning beam in the air without any galvanometer scanning.

In addition to the ERF and the TRF, the flange reference frame (FRF), base reference frame (BRF), and world reference frame (WRF) are also defined. The FRF is associated with the end joint of the robotic arm, which the sample arm of our vis-OCT attaches to. When calibrating the TRF to align with the vis-OCT imaging volume spatially, we specified the pose of the TRF relative to the FRF. The BRF is associated with the base of the robotic arm, and the WRF is the main static reference frame, the position of which is encoded relative to the BRF. Since the position of the TRF relative to the WRF can be read out and defined by the Meca500, we set the WRF as the ERF for convenience.

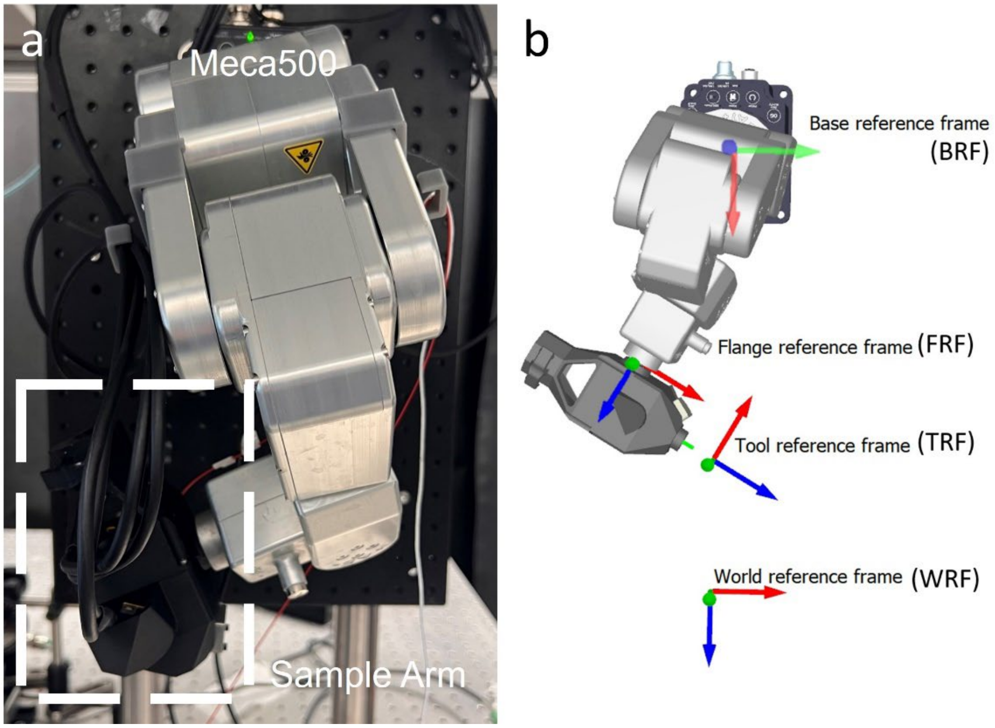


**Fig. S1. Robotic arm and definition of reference frames**. a) Sample arm of freeform robotic vis-OCT, with sample arm enclosed in a white dashed square. b) Illustration of several of the reference frames associated with the robotic arm. The FRF is the reference frame associated with the end joint of the robotic arm and the BRF is the frame associated with the base of the robotic arm. The TRF is associated with the OCT volumetric image and WRF is associated with the object being imaged. The pose of the TRF relative to the FRF and the pose of the WRF to the BRF are user-defined.

**Supplemental Methods 1: Calibration of Tool Reference Frame**

We calibrated the TRF to coordinate the TRF and the OCT imaging volume coordinate systems. First, we spatially aligned the TCP, which is the origin of the TRF, with the focal point of our vis-OCT system without galvanometer scanning. Overall, this step aligns the TCP with the origin of the OCT imaging volume. We aligned the origins by imaging a stationary target point at the focal plane and the center of the OCT FOV at multiple robot arm orientations (Fig. S2a). Following the collection of the robot joint angles used to image the target point, we utilized the "Define TCP" tool of the commercial RoboDK software to determine the point the robot arm was rotating around, which is the focal point of the robotic vis-OCT. To ensure that the stationary target point was located at the vis-OCT focal point, we adjusted the sample arm position to maximize the OCT signal returning from a mirror without galvanometer scanning and set the target point to have the same position as the mirror.

To calibrate the TCP, we set the corner of a fixed rectangle on a plastic USAF51 target card attached to the top of a piece of paper as our stationary target point (Fig. S2b). We adjusted the robot arm posture until the corner was at the lateral center of the vis-OCT FOV (intersection of blue and red lines in Fig. S2b). Next, we moved the robot arm so that the surface corner at the center of the FOV (vertical red line in Fig. 2c) was located at the axial position corresponding to the depth of the OCT focal plane (dotted green line in Fig. S2c). We repeated this process several times to obtain multiple orientations to view the corner of the stationary target point at the focal point of the vis-OCT. After recording four different poses, we inputted the joint positions into the "Define TCP" tool of the RoboDK software. The "Define TCP" tool determines the spatial location of the point about which the robot arm rotates, which is the OCT focal point. After the initial measurement of the TCP, we added poses and recalculated the TCP. Additional poses were added until the TCP position converged, with the TCP shifting by less than 0.1 mm with the addition of another joint position. Overall, we used a total of 6 points before convergence of the TCP.

To orient the z-axis of the TRF along the primary optical axis of the OCT and the x and y-axes along the scanning directions of the galvanometer, we minimized deviations between the relative angular orientation of the OCT image volume and the current TRF. An initial estimate for the relative pose of the OCT image volume to the FRF was taken based on the mechanical design of the sample arm, and the relative orientation of the TRF to the FRF was set according to this estimate. For a specified cartesian axis direction, we translated the robotic arm position in the TRF by 500 μm while imaging the stationary target point. After each translation, we measured the difference in the target point position within the OCT imaging volume by comparing the target point position within the OCT imaging volume before and after the movement (Fig. S2d). After each measurement, we changed the orientation of the TRF relative to the FRF in increments of 0.5 degrees along the axes perpendicular to the movement axis until the off-axis distance the target point moved after the translation was minimized. Once the distance is minimized, the relative orientation between the OCT image volume and TRF along the specified axis is optimized. We repeated this process for all axes.

To assess the accuracy of our TCP calibration, we positioned our USAF51 corner reference point at the focal point of our vis-OCT and rotated our sample arm around that point. Since the TRF rotates about the TCP, the reference point should remain in the same position in the OCT volume regardless of rotation if the TCP was calibrated correctly to the focal point. Thus, we placed our USAF51 corner reference point at the focal point and rotated our TRF +10 and -10 degrees about the x, y, and z-axis of the TRF in increments of 2 degrees. We found that the TCP shifted by less than 200 μm from the initial position in the imaging volume for every rotational orientation (Fig. S2e), which is within the positional accuracy of the robot arm[2]. Additionally, tension from the galvanometer cable may also contribute to positional errors following rotations.

To access the calibration of axes directions, we translated our robot arm 200 μm in the +z, -z, +y, -y, +x, and -x directions and measured off-axis drifts in the OCT imaging volume. Following each translation, we recorded the spatial coordinates of our reference point in the OCT imaging volume relative to the original position with no translation. The off-axis translation, defined as translational changes in the reference point location along the axes that were not translated in the TRF but translated in the OCT image plane, was 13.3 ± 3.8 μm (Fig. S2f). This off-axis deviation was not statistically significant from the lateral resolution of our OCT, which was about 8 μm.

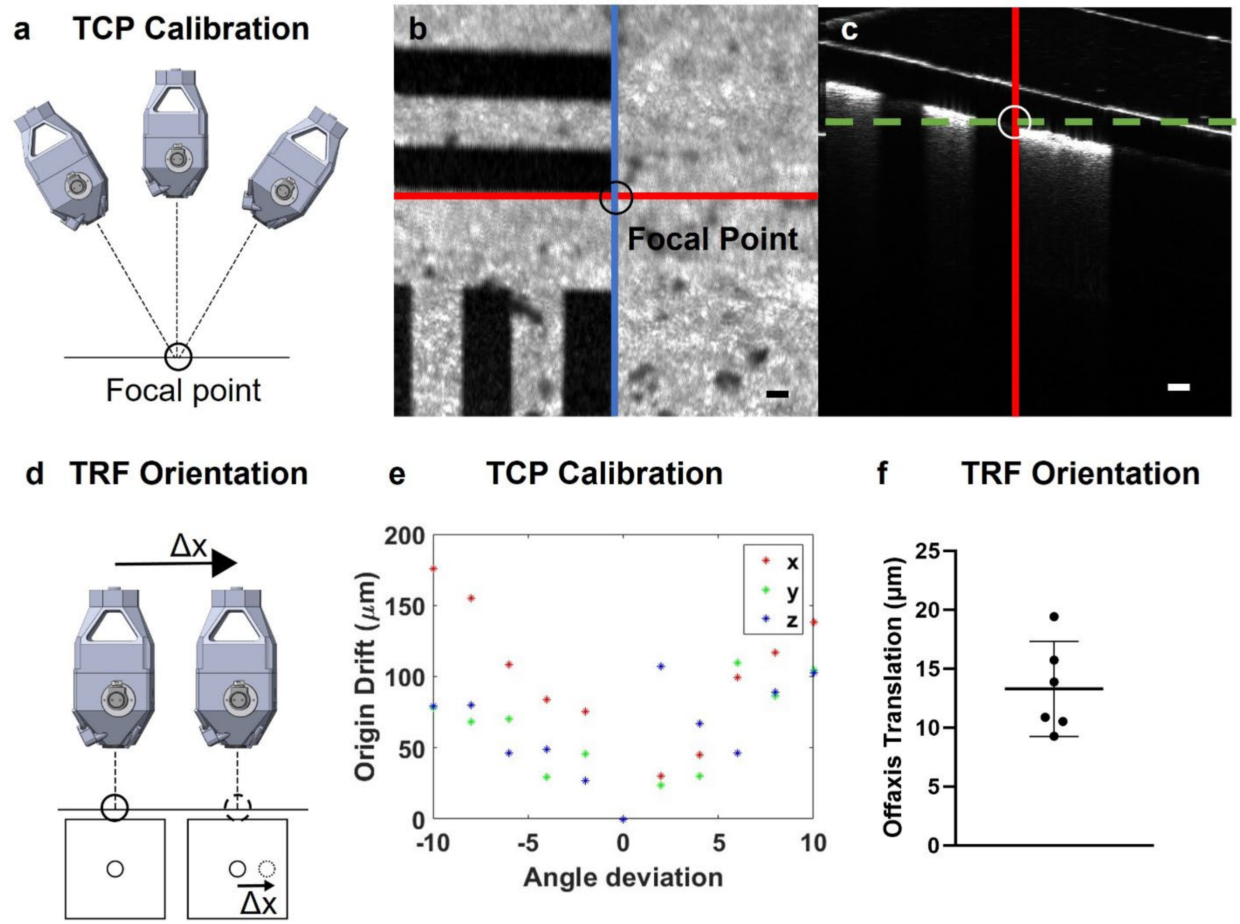


**Fig S2. Calibration of tool reference frame.** a) The TCP was calibrated by imaging a target point at the focal point of the OCT volume and recording the robot joint angles at which the target point was at the focal point. The rotational center was the TCP. b) En-face image of a USAF51 target card zoomed into group 1, element 2, which served as our target point. The red and blue lines intersect the lateral position of the reference corner and are at the lateral center of the OCT field of view. c) B-scan of USAF51 card corresponding to the position of the blue line in (a). The red vertical line corresponds to the lateral position of the red line in (a). The dotted green line corresponds to the axial depth of the target card at the corner point. The scale bars are 100 μm. d) We calibrated the TRF orientation by minimizing off-axis translations following movement in the TRF. e) We assessed TCP calibration accuracy by measuring the translational drift of the reference point from the initial position as the robotic arm rotates about the TRF. Drifts in the origin are within the robot arm's position accuracy. f) Off-axis translational shifts as the robotic arm moved 200 μm in the TRF in each cartesian direction to assess TRF orientation. Drifts are not statistically different than the lateral resolution of vis-OCT.

**Supplemental Methods 2: Image Acquisition Process for Outflow Pathway**

We rotated the TRF about the Z-axis of the ERF to image the entire AHO pathway. Thus, to determine a trajectory for the robotic OCT around the limbus of the eye, the crucial step is to determine the pose of the ERF coordinate system relative to the TRF (Fig. S3a). To accomplish this, we (1) identify the optical axis of the eye, (2) estimate the origin of the ERF, and (3) acquire data scanning 360 degrees around the outflow pathway (Fig. S3b). More details of this procedure are given below.

**Step 1**: We brought the central cornea of the mouse into the OCT imaging volume FOV. After the central cornea was brought into the FOV, we acquired real-time B-scans of the cornea along the x and y-axes of the TRF for subsequent image analysis.

- We usually did this step manually by centering the eye in the pupil camera's FOV, although it is possible to do this step automatically. A method to do this automatically with the pupil camera is to identify the position of the pupil in the pupil camera, calculate the offset of the pupil position to the center of the FOV, and translate the TRF by the offset. The TOF sensor on the sample arm can then be used to translate the TRF z-axis origin until the mouse cornea is focused, with the eye 25mm away from the sample arm, which is the focal length of our vis-OCT.

**Step 2:** We parallelized the z-axis of the TRF and the Z-axis of the ERF (aligning the pose of the TRF such that R and P angles are zero relative to the ERF). This is equal to aligning the pose of the TRF such that the xy-plane of the TRF is parallel to the XY-plane of the ERF. For input data, we analyzed real-time B-scans along the xz-plane and yz-plane in the TRF. Using the hypothesis that the primary symmetry axis within each of the cornea B-scan images closely resembles the Z-axis in ERF, we sought to parallelize the symmetry axis in each B-scan with the optical axis of the TRF. This was accomplished with the following steps:

- We evaluated the relative orientation of the z-axis of the TRF and the Z-axis of the ERF along a given axis using the slope of a line fit to the center of mass in the axial direction as a function of transverse position x. We calculate the center of mass as a function of position x with the following equation where $I(z,x)$ is the intensity of the central B-scan at depth z and position x:

$$COM(x) = \frac{\Sigma_z I(z,x) z}{\Sigma_z I(z,x)}. \qquad \text{Eq. S1}$$

- After $COM(x)$ was calculated, we fit a line to $COM(x)$ and used the fitted slope as an estimate for relative alignment between the z-axis of the TRF and the Z-axis of the ERF. When the z-axes of the TRF and ERF are well aligned, the fitted slope has a value near zero. The greater the misalignment between the z-axes, the higher the absolute value of the fitted slope, with the sign of the slope indicating whether the z-axis of the TRF was aligned clockwise or counterclockwise to the Z-axis of the ERF along the scan axis which the B-scan was acquired.

- Based on the slope of a line fit to $COM(x)$, we rotated the sample arm around the scanning axis in the TRF by angle c*slope($COM(x)$), where c is a constant influencing the speed of the alignment process. We rotated the x-axis of the TRF based on the fast-scanning axis B-scan and the y-axis for the slow-scanning axis B-scan. Overall, this adjustment changes the R and P angle of the TRF pose relative to the ERF. We continued this process until we were satisfied with the relative orientation of the TRF z-axis and ERF Z-axis.
- Although $COM(x)$ is curved and not strictly linear, we found the above method to be simple and robust for evaluating the relative orientation of the TRF z-axis and ERF Z-axis when tested on a mouse eye and anterior segment eye phantom. Overall, we note that the slope fitting method performed equally or more effectively as compared to using the relative position of the iris on both sides of the B-scan and evaluating the tilt of the lens. Moreover, the step of fitting a line to $COM(x)$ could be considered approximating $COM_y(x)$ with a first-order Taylor approximation. We found a high correlation between the value of the slope of $COM(x)$ with the slope of manual lines drawn parallel to the optical axis of the eye (correlation of .967).
- Fig. S3c illustrates the process of parallelizing the z-axis of the TRF and the Z-axis of the ERF. The red and blue shaded sample arms in the top panel correspond to poses where the z-axis of the TRF is parallel and oblique to the Z-axis of the ERF, respectively. Representative B-scans, where the z-axis of the TRF and the Z-axis are parallel and oblique to each other, are shown underneath the red and blue boxes.

**<u>Step 3:</u>** We spatially overlapped the z-axis of the TRF with the Z-axis in ERF. This process aligns the x and y origin of the TRF with the X and Y origin of the ERF. To determine the X and Y origin of the ERF, we located the apex of the cornea in the B-scans (see supplemental methods 3) and translated the TRF to move the apex into the center of the OCT FOV.

- We used the lateral position of the apex of the fitted cornea in the central B-scans as an estimate of the origin of the ERF along the galvanometer scanning directions. Based on the offset between the apex and central A-line, we translated the TRF by the corresponding spatial distance. After laterally moving the TRF x and y origins to the lateral position of the apex of the cornea, the position of the real-time segmented cornea was 1.5 mm (roughly the radius of our mouse eyes) above the origin of the ERF in the Z-axis of the ERF.

- Fig. S3d illustrates the process of identifying the origin of the X and Y-axes of the ERF. The sample arm shaded in red shows a position where the origin of the TRF is spatially offset from the origin of the ERF in the lateral direction. The position of the eye relative to the sample arm and the central B-scan corresponding to the offset sample arm position are shown below in the red boxes. In the red boxes, the pupil is offset to the left from the center of FOV and the top of the cornea is offset to the left from the middle a-line of the central B-scan. The sample arm shaded in blue corresponds to a position where the lateral origins of the TRF and ERF coincide. The position of the eye and corresponding central B-scan are found below in the blue boxes. In the blue boxes, the pupil is in the center of the FOV and the apex of the cornea is in the center of the central B-scan.

**Step 4:** We determined the origin of the ERF along the Z-axis by setting the origin of the ERF 1.5 mm underneath the apex of the cornea along the Z-axis of the ERF (found in the previous step).

- We repeated steps 2 and 3 if 1) the z-axis of the TRF and Z-axis of the ERF were not parallel or 2) the x and y origin of the TRF were not well aligned with the apex of the cornea.

**Step 5:** We determined the remaining DoF of RPW angle (orientation of X, Y-axis within the defined XY-plane in ERF) by recording the relative position of eye quadrants to the galvanometer scanning axes.

- After the alignment of the z-axis of the TRF and the Z-axis of the ERF, the XY-axes of the ERF are oriented in the same direction as the current xy-axes of the TRF after step 3. Thus, we recorded the orientation of each quadrant relative to the galvanometer scanning axes before image acquisition. Before each imaging session, we set the default position of the TRF to a default pose relative to the robot base, with the scanning axes having a fixed orientation relative to the edges of the optical table. The xy-axes of the TRF undergo rotations during steps 2 and 3.

**Step 6:** We mapped out a trajectory for rotating the TRF around the ERF.

- Following the definition of ERF, we set the WRF as the ERF.
- Next, we performed conformal scanning of the outflow pathway to acquire data.

- The TRF was rotated 60 degrees around the X-axis of the ERF. Empirically, the outflow pathway is approximately 60 degrees from the Z-axis of the ERF, meaning that the scanning conformal to the outflow pathway.
- The TRF is rotated 360 degrees around the Z-axis of the ERF, with minor adjustments in position made every 45 degrees to optimize signal quality. After every 45 degrees, we captured an OCT volume for a total of eight volumes. Several of the minor adjustments we made include translating the TRF until the surface topology of the eye was symmetrical about the x-axis of the TRF and moving SC into the central A-line in the central B-scan. Additionally, we rotated around the TRF xy-axes when the optical axis of the OCT was not conformal to the surface, although this was usually not the case. For conformal imaging, the minor axis of an ellipse fit to SC should orient along the z-axis of the TRF.
- In cases where we were unsatisfied with the ERF origin position, we calibrated the ERF origin using an alternative method. For this calibration, we imaged the outflow pathway at a conformal angle at 4 positions around the eye. We inputted the robot joint angles at these 4 positions as well as during the original alignment position after step 3 to the “Define TCP” tool of a commercial robot motion simulation software, RoboDK to refine the origin of the ERF.

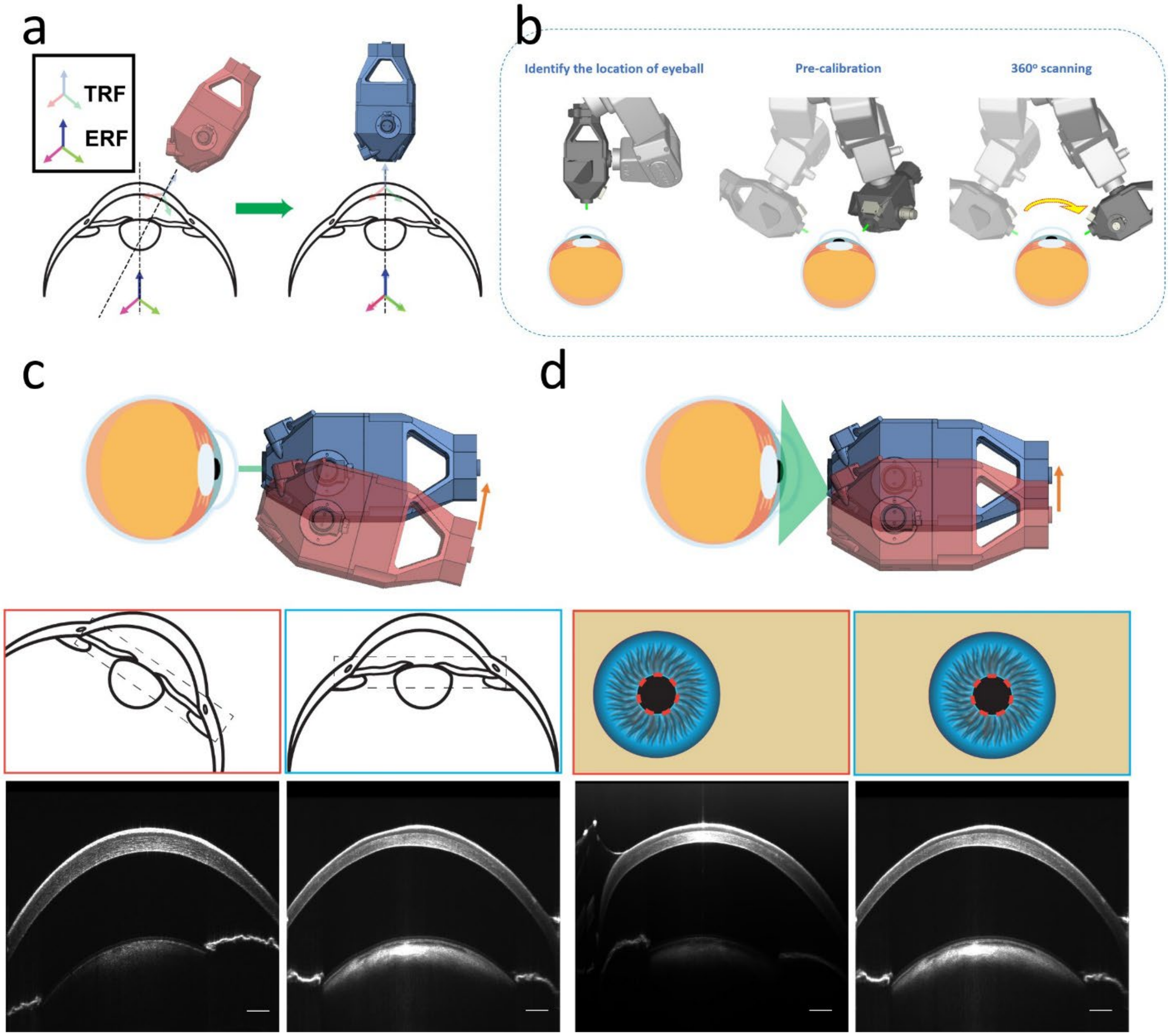


**Fig S3. Image Acquisition Protocol for Outflow Pathway**. a) Overview of the calibration process for imaging the limbal region of the eye, which equates to determining the pose of the TRF relative to the TRF. b) Overview of the data acquisition scheme, which involves identifying the eye, aligning the Z-axis of the ERF and z-axis of the TRF, aligning the lateral origins of the ERF and TRF, and rotating the sample arm around the ERF Z-axis. c) Real-time OCT images are used to align the z-axes of the TRF are ERF. Optimized positioning is shown in blue, and suboptimal positioning is shown in red. d) The pupil camera and real-time OCT images are used to align the lateral origins of the ERF and TRF. Optimized positioning is shown in blue, and suboptimal positioning is shown in red. Scale bars are 200 µm.

**Supplemental Methods 3: Real-time corneal apex segmentation**

We performed real-time image processing of the central B-scans for both scanning axes to locate the origin of the ERF along its X and Y-axes. First, we acquired an initial raw image (Fig. S4a) and then mean-filtered (Fig. S4b) the image. The original image is sized 1024x128 and we used a mean filter of size 9x3. For faster processing, we downsampled the image by a factor of 8 in the axial direction, at which point the preview image was sized 128x128. We automatically segmented the cornea and lens using a modified hysteresis thresholding scheme. Overall, we found hysteresis thresholding effective for getting rid of auto-correlation near the zero-delay line when an unbalanced preview was used. For a balanced preview, thresholding without any additional steps was used. For hysteresis thresholding, we binarized the image with an automatically defined high threshold ($85^{th}$ percentile of image intensity, Fig. S4c) and removed regions less than 20 percent of the size of the largest connected binarized region. Next, we binarized the image to include pixels with intensity greater than a low threshold ($65^{th}$ percentile of image intensity, Fig. S4d) but less than the high threshold ($85^{th}$ percentile). For pixels with intensity greater than the low threshold but less than the high threshold, we considered any term connected to the border as an autocorrelation signal. We kept any signal with intensity greater than the low threshold but not autocorrelation as the structural mask (Fig. S4e). We removed all subregions in the structural mask with an area less than 20 percent of the largest region. We fit a third-degree polynomial to the top border of the binarized structural mask (Fig. S4f) to identify the top surface of the cornea. We used the lateral position of the apex of the fitted cornea to estimate the origin of the ERF along the scanning direction.

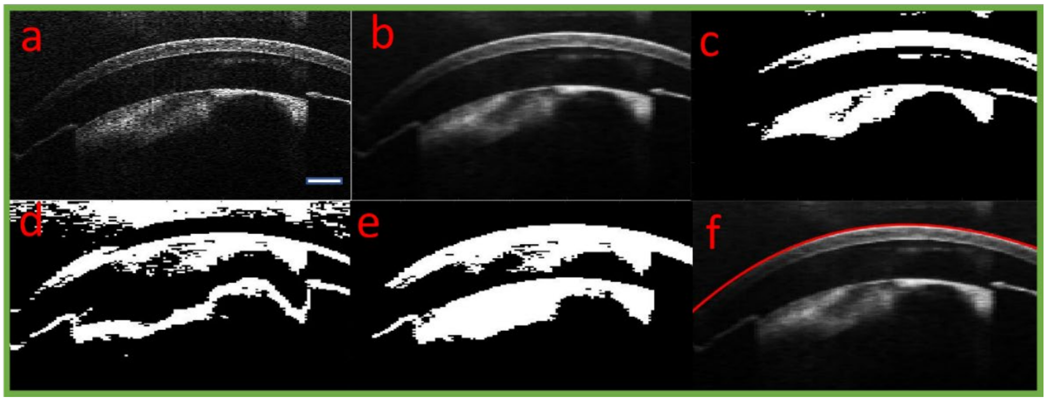


**Fig. S4. Real-time image processing to segment cornea**. a) Raw B-scan image in real-time. b) Box-filtered image. c) Image in (b) binarized with a high threshold. d) Image in (b) binarized with intensity greater than low threshold but less than high threshold. e) Combination of (c) and (d) minus components touching boundaries. f) Raw image with the top of the cornea overlayed in red. The scale bar is 200 μm.

**Supplemental Methods 4: Simultaneous Localization and Mapping for OCT Volumetric Montaging**

We developed a volumetric registration algorithm based on simultaneous localization and mapping (SLAM) to montage the volumetric OCT data acquired from different poses. Commonly used in robotics, SLAM is the process of mapping out an environment while keeping track of an object's location within that environment [4]. Typically, a combination of the robot's internally measured position and external sensor readings are used to localize the robot's position within the environment. Since robots have limited positional accuracy, algorithms often integrate external sensor data to correct the robot's internally measured position. In our case, we measured the location of the robot relative to the eye by reading the pose of the TRF relative to the ERF for each acquired volume. To correct errors in this reading, we matched anatomical structures as landmark points in the overlapping regions between volumes. In other words, the acquired volumes served as the external sensors used to correct the position reading of our robotic arm. We detail the steps of our OCT volumetric merging below (Fig. S5a).

**Segmentation of Eye Surface and Representation as Point Cloud**

For each volume, we represented the outer surface of the eye closest to the incident OCT beam as a point cloud. We define $I(x, y, z)$ as the intensity of the OCT volume at coordinate (x,y,z) in the acquired TRF reference frame (Fig. S5b). First, we binarized each B-scan by thresholding (using the $10^{th}$ percentile of intensity as cutoff) each B-scan and only keeping the largest spatially connected remaining object (Fig. S5c). We took the union of all the binarized B-scans to form a binarized volume.

After segmentation, we observed a continuous corneal and limbal structure (Fig. S5d). For every A-line in the binarized volume, we took the axial position z at the outer surface of the segmented volume and represented this set of points as a point cloud (Fig. S5e).

**Montaging of Overlapping Point Clouds**

To determine whether two volumes A and B overlap, we checked whether the imaging volumes of A and B had greater than 20% volumetric overlap in the ERF. To determine the imaging volume of volume A in the ERF, we applied the relative pose of the TRF to the ERF when A was acquired to the initial imaging volume of A in its reference frame. The initial imaging volume of A in its reference frame was a cuboid with a range of (-imaging range/2, imaging range/2) in the x and y axes and a range of (0, $z_{max}$) for the z-axis, where

$$z_{max} = \frac{\pi}{2\delta k} \quad \text{Eq. S2}$$

and $\delta k$ is the wavenumber spacing between pixels for the spectrometer. Similarly, we determined the imaging volume of B in the ERF. If the volume of the intersecting regions of the imaging volumes A and B in the ERF had a volume greater than 20% that of volumes A and B, we considered volumes A and B to be overlapping. If A and B had sufficient overlap, we merged the two point clouds of A and B with the following procedure.

Let $P_A$ be the point cloud for the outer surface of volume A. Furthermore, let $S_A$ and $S_B$ be the functions returning the axial position of the eye surface for two overlapping volumes A and B. For an arbitrary surface N, if the outer surface of the segmented surface at $(x_i, y_j)$ is $z_{ij}$, then

$$S_N(x_i, y_j) = z_{ij}. \quad \text{Eq S3}$$

We identified the lateral position of 15 different common landmark points, usually vessel branch points in the OCTA, in the overlapping region between A and B (Fig. S5f-g). Supposing that the landmark points have lateral positions $\{(x_1, y_1), \ldots, (x_{15}, y_{15})\}$ in volume A in the ERF, then the set of landmark points $L_A$ is defined as

$$L_A = \{(x_1, y_1, S_A(x_1, y_1)), \ldots, (x_{15}, y_{15}, S_A(x_{15}, y_{15}))\}. \quad \text{Eq. S4}$$

The landmark points $L_B$ in volume B are defined in the same fashion. We applied the M-estimator sample consensus algorithm to determine the rigid transformation $T_{AB}$ mapping $L_A$ onto $L_B$ (Fig. S5h)[5].

After calculating the rigid transformations mapping each imaging volume with its overlapping neighboring volumes, we found the transformation mapping each imaging volume to the ERF. First, we choose a volume to serve as an initial stationary volume. For purposes of discussion, let volume A be the stationary volume and $T_{A,ERF}$ be the relative pose of the TRF to the ERF when volume A was acquired. For volume A, the transformation mapping A to the ERF is $T_{A,ERF}$. Since rigid transformations are linear operators, the rigid transformation mapping volumes to each other can be applied in sequence to map all volumes onto the same coordinate system. If $T_{B,A}$ maps volume B to volume A, then $T_{B,A} * T_{A,ERF}$ would map volume B to the ERF.

After determining initial rigid transformations mapping each volume to the ERF, we applied the transformations to each point cloud. Next, we used the M-estimator Sample Consensus algorithm to fit a sphere to the union of all the volumetric point clouds, to find the centroid of the fitted sphere, and to determine the inlier points used for the sphere fitting. Using the inlier points, we found a rotational axis of symmetry representing the visual axis of the eye. After determining the centroid of the fitted sphere and the visual axis of the eye, we updated each of the initial transformations. Specifically, if $T_{Origin}$ translates the origin of the fitted sphere to the origin of the ERF and $T_{Axis}$ aligns the calculated visual axis with the current visual axis of the ERF, we updated the rigid transformation for volume A to the ERF as

$$T_{A,ERF} = T_{Axis}T_{Origin}T_{A,ERF}. \qquad \text{Eq. S5}$$

We also updated the transformations for all the other volumes in the same way.

**Loop Closure Problem**

When a series of transformations are applied consecutively, errors from each transformation cascade. In our case, this may result in a mismatch between surfaces if a volume is overlapping with multiple different volumes.

For each pair of point clouds $P_A$ and $P_B$ that were montaged together, we determined the points in the overlapping regions between $P_A$ and $P_B$ (Fig. S5i). After applying $T_{A,B}$ to $P_A$, $T_{A,B}(P_A)$ and $P_B$ reside in the same coordinate system. For each point in $T_{A,B}(P_A)$, we found the closest point in $P_B$. We considered the union of closest points in $P_B$ to $T_{A,B}(P_A)$ as the overlapping region in $P_B$, which we denote $O_{BA}$. We found the overlapping region $O_{AB}$ in the same manner. We define $O_{ERF}$ as the union of all overlapping regions between every overlapping volume. After determining all sets of overlapping regions in the ERF, we refined the transformation mapping each volume to the ERF.

To refine the transformation mapping volume B to the ERF, we determined the transformation mapping the overlapping regions of B to the union of all overlapping regions. Supposing volume B overlapped with volumes A and C, we found the set of overlapping regions $O_{BA}$ and $O_{BC}$. Then, we utilized the iterative closest point algorithm to calculate the transformation $T_{OB,OERF}$ mapping $O_{BA}$ and $O_{BC}$ onto $O_{ERF} - (O_{BA}\ U\ O_{BC})$[6]. If $T_{B,ERF}$ initially mapped volume B to the ERF, then we updated the following:

$$T_{B,ERF} = T_{OB,OERF} * T_{B,ERF}, \text{ and} \qquad \text{Eq. S6}$$

$$O_{BA} = T_{OB,OERF}(O_{BA}), O_{BC} = T_{OB,OERF}(O_{BC}). \qquad \text{Eq. S7}$$

After each step, we updated $O_{ERF}$ to be the union of all overlapping regions using the updated overlapping regions found above. We applied this process to every volume consecutively to refine the transformation of all volumes to the ERF (Fig. S5j). After refining every transformation, we repeated the process of refining every transformation two more times. Following transformation refinement, the union of all point clouds mapped onto the ERF is smooth and continuous (Fig. S5k).

**Volumetric Montaging**

After identifying the rigid transformations $T_A$ mapping $P_A$ from its original image reference frame to the TRF for every volume, we applied the transformation to every volume. If $T_A$ maps point (x,y,z) from the original image coordinate system of A to point (i,j,k) in the ERF, then:

$$I_{world}(i,j,k) = I_A(x,y,z). \quad \text{Eq. S8}$$

When multiple points from different volumes are mapped onto the same point in the ERF, we took the maximum intensity of the volumes mapping onto that point. In other words, if $T_A$ maps $(x_a, y_a, z_a)$ to (i,j,k) and $T_B$ maps $(x_b, y_b, z_b)$ to (i,j,k), then $I_{world}(i,j,k)$ would equate to the maximum of $I_A(x_a, y_a, z_a)$ and $I_B(x_b, y_b, z_b)$.

**Representation of Data within Spherical Coordinates**

We transformed the reconstructed 3D volume from cartesian to spherical coordinates using the following transformation

$$I_{spherical}(r,\theta,\phi) = I_{world}(rsin\theta cos\phi, rsin\theta sin\phi, rcos\theta). \quad \text{Eq S9}$$

En-face projections E were created as a function of $\theta$ and $\phi$ by

$$E(\theta,\phi) = \frac{\Sigma_{r=r_-}^{r=r^+} I_{sperical}(r,\theta,\phi)}{\Sigma_{r=r_-}^{r=r^+} 1}. \quad \text{Eq. S10}$$

As a spherical surface cannot be perfectly projected onto a plane, we applied the various projections from cartography to $E(\theta,\phi)$ to render the eye in 2D (Fig. S5l)[7].

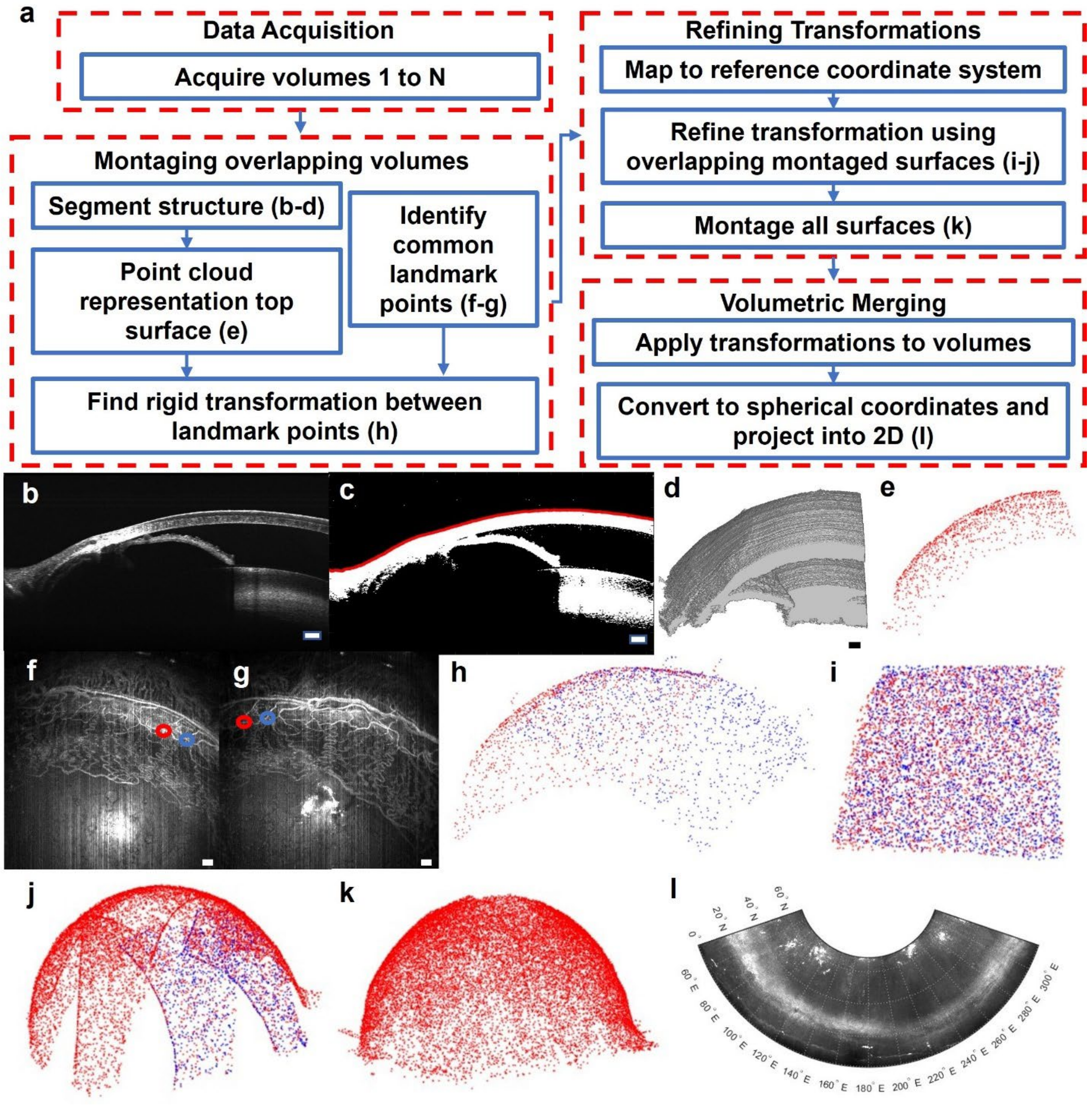


**Fig S5. Volumetric registration of different OCT volumes**. a) Summary of steps. b) Acquired B-scan. c) B-scan thresholded and binarized, with the outer eye surface marked in red. d) 3D reconstruction of the binarized structure. e) The outer eye surface is represented as a point cloud. f + g) OCTA of overlapping regions. Two pairs of common vessel branch points between the OCTA are marked by blue and red circles. h) Montaging of adjacent surface point clouds. The point cloud of the first volume is shown in red and the point cloud of the second volume is shown in blue. i) Montaging of adjacent surface point clouds in the overlapping regions. j) The overlapping region of one point cloud is shown in blue, with all overlapping regions of all point clouds shown in red. k) Point cloud representation of the entire eye surface. l) Equaconic projection of the eye, with positions along the eye represented as latitude and longitude lines. All point clouds were downsampled by 100 times for visualization purposes. Scale bars are 100 µm.

**Supplemental Methods 5: Reconstruction of Outflow Pathway**

Following volumetric montaging, we reconstructed the AHO pathways. To better visualize the pathway, we projected the structure of the pathway into two dimensions. Below, we detail the methodology used to segment and project the outflow pathway.

**SC and CC segmentation**

To reconstruct the entire SC, we segmented SC for every B-scan in each acquired volume (Fig. S6a). First, we manually selected a point contained within SC every 30 B-scans. For each of these B-scans, we used a flood-fill algorithm to determine the borders of SC. For this algorithm, we found the set of continuous pixels with grayscale intensity below the intensity of the chosen point plus a tolerance value.

Following the semi-automated segmentation of SC every 30 B-scans, we determined the centroid of the SC in each of those B-scans. Based on the SC centroid position, we interpolated the SC centroid position to the scans that had not yet been segmented. Next, we used the same flood-fill algorithm as before to determine the borders of SC for all B-scans. Manual correction to the segmentation was applied as needed. All CCs were manually segmented.

**Resampling to reconstruct SC along its height dimension**

We define G as the 3D volumetric segmentation of SC. The binarized projection mask of SC within each volume is the set of points (x,y) satisfying

$$\Sigma_{\mathrm{z}}\, G(x, y, z) > 0. \qquad \text{Eq. S11}$$

Overall, the binarized projection mask forms a continuous structure, reflecting the connectivity of SC (blue overlay in Fig. S6b). We skeletonized the binarized mask of SC and pruned all branches except for the largest one. Based on the coordinates of the skeleton, we resampled SC along the skeleton, with lateral averaging occurring along the normal direction of the skeleton to improve SNR. Specifically, the volumetric data along the skeleton centerline and 5 μm along the normal direction of the skeleton were averaged together (Fig. S6c). Letting X and Y be the ordered set of points along the skeleton, we define the normal direction of the skeleton as

$$\nabla(\text{skeleton}) = (-\partial Y, \partial X)\,. \qquad \text{Eq. S12}$$

In this case, we computed $\partial Y$ and $\partial X$ numerically calculated by taking the central difference of X and Y. Each A-line of the resampled image corresponds to the A-line at a lateral position of the skeleton in the original volume. Following the resampling of SC, we flattened the top surface of the eye to have the same depth position in the resampled image (Fig. S6d). Flattening was done by thresholding the resampled image, taking the largest continuous region, and translating each A-line such that the top border of the segmented image ended up at the same vertical depth position. To stitch the resampled SC between overlapping volumes, we determined the positions where SC overlapped in the global ERF coordinate system after SLAM. The skeleton points in the overlapping region were determined and matched for both overlapping volumes. For each matched skeleton point, we resample at the position closer to the center of the original single-volume field of view.

**Resampling to reconstruct SC along its width dimension**

For the set of points in the binarized mask of SC, we found the mean depth of SC along each position $z_{sc}(x,y)$

$$z_{sc}(x,y) = \frac{\Sigma_z G(x,y,z)z}{\Sigma_z G(x,y,z)}. \quad \text{Eq. S13}$$

The intensity of the resampled projected image at P is

$$P(x,y) = I_{world}(x,y,z_{SC}(x,y)). \quad \text{Eq. S14}$$

For lateral positions not contained in the binarized mask of SC, where $\Sigma_z G(x,y,z) = 0$, we interpolated $z_{SC}(x,y)$ at these points using a nearest neighbor approach to find the depth to resample at.

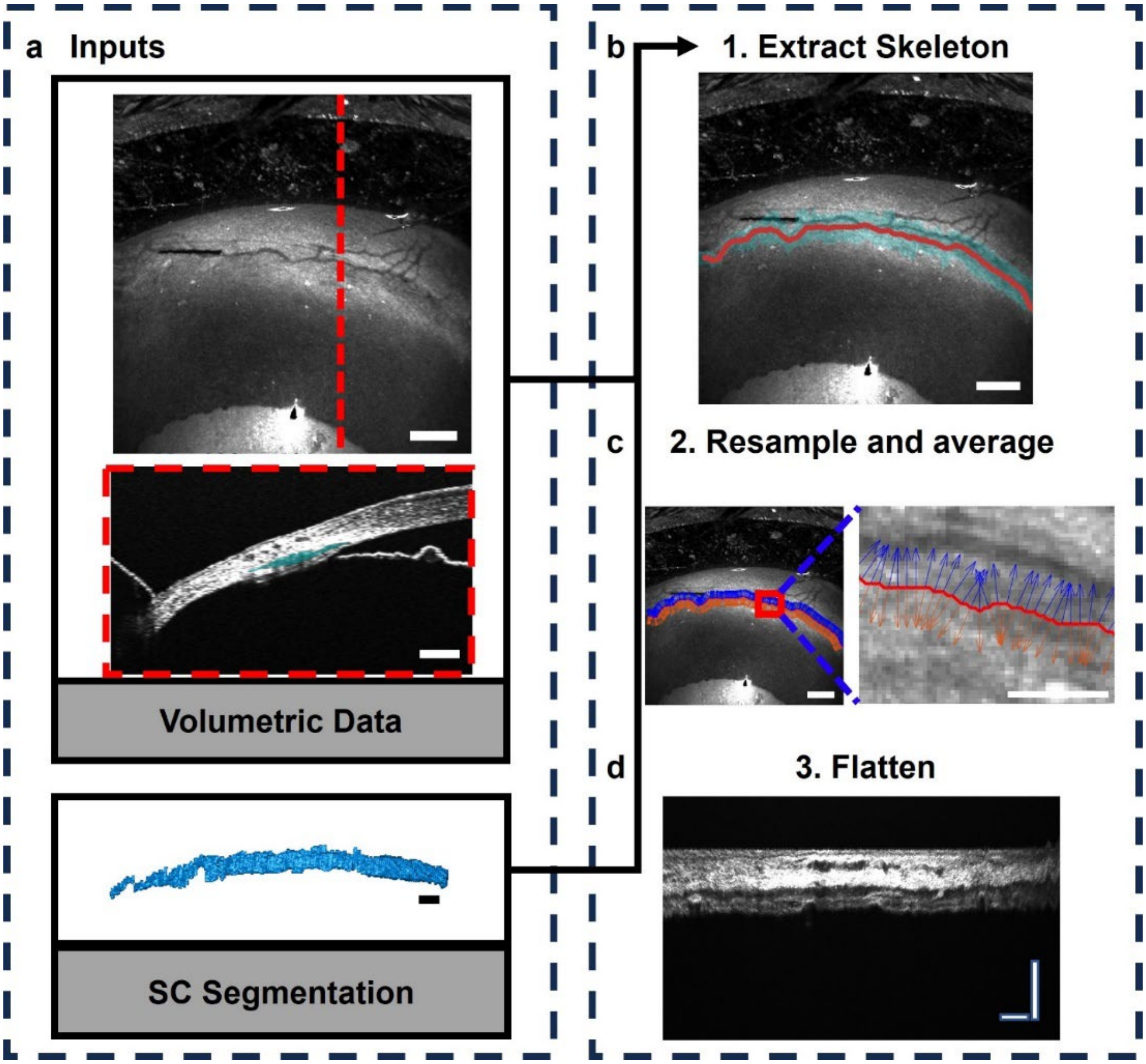


**Fig S6. Resampling of Schlemm's Canal**. a) Input data used to resample SC in the height dimension for a given volume. Volumetric OCT data and volumetric segmentation of SC are inputs to the resampling data. An OCT en face projection with the B-scan at the position of the vertical dotted red line is shown in the volumetric data panel. The SC is shaded in blue in the B-scan. The scale bar for the OCT en face is 200 µm and the scale bars for the B-scan and SC segmentation are 100 µm. b) We determined the skeleton along the lateral positions of the segmented SC. The skeleton is shown in the orange line and the lateral positions of the SC are shaded in light blue. The scale bar is 200 µm. c) For the skeleton, we determined the normal direction of the skeleton along each point in the skeleton. The red line shows the skeleton and the orange and blue lines show the normal vectors of the skeleton. The image to the right is a zoomed-in version of the red box in the left image. The scale bar on the left is 200 µm and the scale bar on the right is 50 µm. d) SC resampled along the skeleton with averaging along the normal direction to the skeleton. The top boundary of each B-scan was flattened to the same axial position. The scale bars are 100 µm in each direction.

**Graph theory for the reconstruction of collector channels**

Given a segmentation of each CC and the volumetric data, we projected each CC connecting SC to the distal vasculature. Each CC was manually segmented. We define C as a set of voxels in the 3D segmentation for an individual CC and the adjacent SC to the CC (Fig. S7a). We modeled C as a graph, where every voxel in C is represented as a node and spatially connected voxels have an edge between them (Fig. S7b)[8]. If the points $(x_i, y_i, z_i)$ and $(x_j, y_j, z_j)$ were spatially connected in C, we assigned the weight of the edge between the nodes associated with points $(x_i, y_i, z_i)$ and $(x_j, y_j, z_j)$ as

$$W((x_i, y_i, z_i), (x_j, y_j, z_j)) = I(x_i, y_i, z_i) + I(x_j, y_j, z_j). \qquad \text{Eq. S15}$$

To determine the end nodes of the graph, we first found the points in C with the lowest depth position relative to the eye surface in the first and last indexed B-scan containing any voxel associated with the nodes in C. We repeated the same process for the point with the highest depth position, after which there were four candidate points for the end nodes of the graph. We set the candidate node with the highest depth dimension as the first end node. In the case that the voxel with the highest depth dimension was in the first indexed B-scan, we assigned the second end node as the voxel with the lowest depth dimension in the last indexed B-scan. Otherwise, we assigned the voxel with the lowest depth dimension in the first indexed B-scan as the second end node. Following the identification of end nodes, we determined the shortest path in the graph between the end nodes using the Dijkstra algorithm (Fig. S7c). The nodes contained within the shortest path were converted to physical voxels within C (Fig. S7d). The lateral position for each of the voxels was taken, with repeated positions removed, as points to resample for the projection of the CC (Fig. S7e). No averaging along the normal direction was done as averaging obscures visualization of CC at its smallest point. After resampling along the lateral positions corresponding to the voxels in the shortest path, a clear lumen connecting SC to the distal vasculature is easily visualized as the resampled CC. (Fig. S7f).

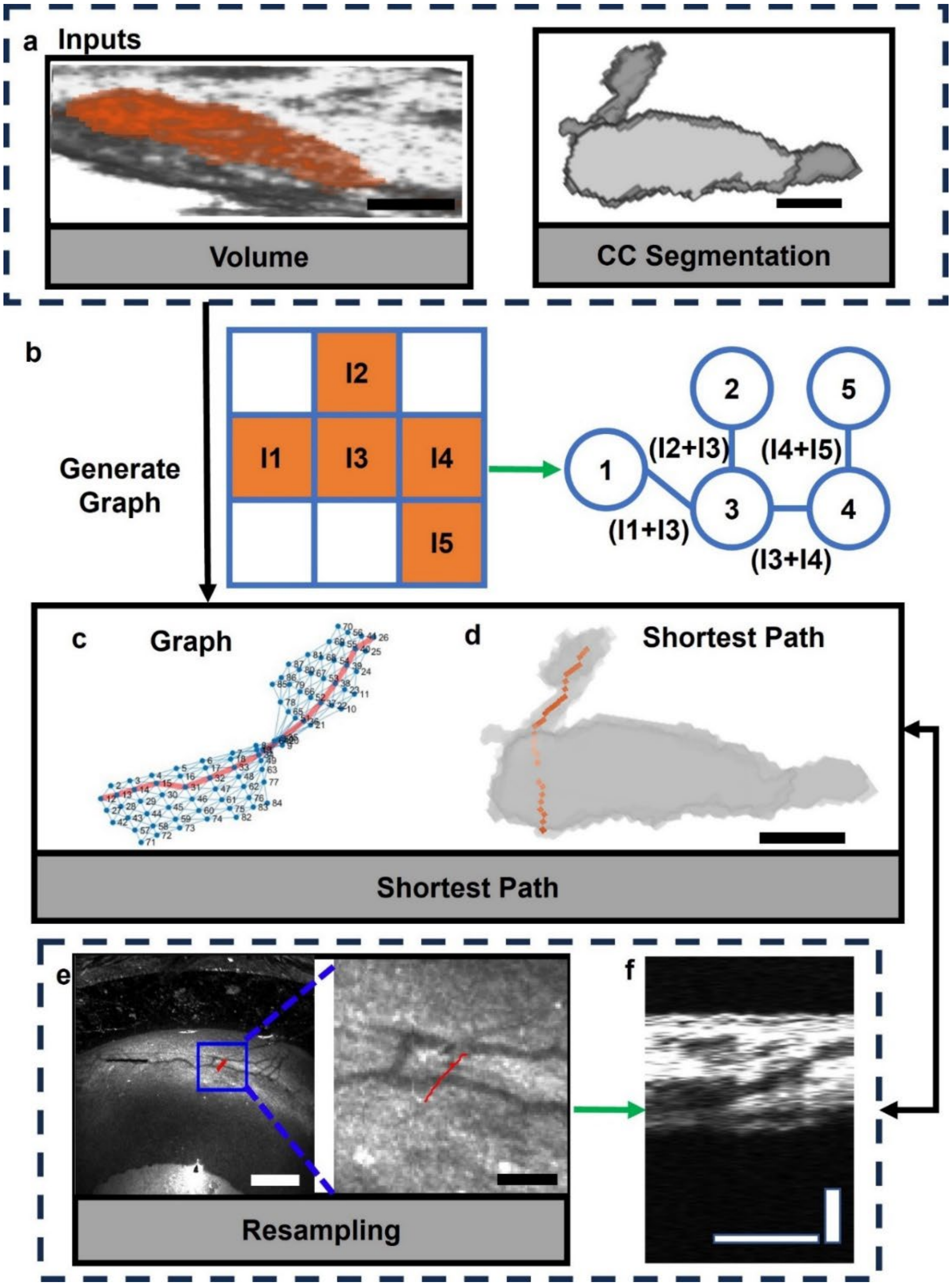


**Fig. S7. Resampling of Collectors Channels**. a) Input data used to resample each CC from SC to distal vasculature. OCT volumetric and each CC segmentation were used as input data for the resampling. The segmentation of CC and adjacent SC in the volume are overlaid in orange. b) The CC segmentation was converted into a graph, with each pixel represented as a node. c) Example graph for a singular B-scan with the shortest path between two nodes outlined in red. d) The pixels corresponding to the shortest path overlaid in orange over the segmented CC and adjacent SC. e) The volumetric data was resampled along the lateral positions of the shortest path (path in red). The image on the right is zoomed in to the blue box on the left. f) CC resampled from SC to distal vasculature using the path shown in red in (e). The scale bars in (a) and (f) are 50 µm. The scale bar in (d) is 100 µm. The left scale bar in (e) is 100 µm and the right scale bar is 40 µm.

**Supplemental Methods 6: Eye Exposure**

As the eyelids typically block the nasal and temporal limbal region (Fig. S8a), we exposed these areas to image the entire limbal region around 360 degrees of the eye. First, we made a small cut to the nasal and temporal portions of the eyelid and inserted a speculum underneath the eyelids (Fig. S8b). An alternative to exposing the eyelid is to proptose the eye and suture the eyelids together underneath the eye (Fig. S8c). However, this process causes blood reflux, significantly enlarging the conventional outflow pathway[9].

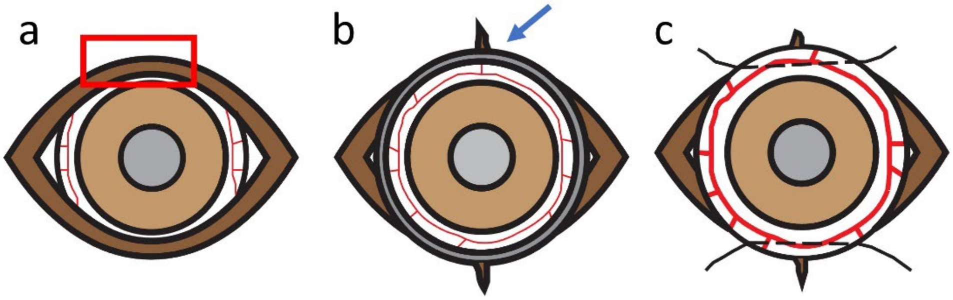


**Fig. S8. Operations done to visualize entire 360 degrees of the outflow pathway**. a) The eyelid (red rectangle) blocks the outflow pathway around the nasal and temporal limbal regions. b) To expose the outflow pathway, small cuts in the temporal and nasal portion of the eye were made and a speculum (blue arrow) was used to expose the outflow pathway. c) As an alternative, after small cuts were made in the temporal and nasal portion of the eye, the eye was proptosed by placing sutures underneath the globe. However, proptosis of the eye causes mild blood reflux.

**Supplemental Methods 7: Real-time image processing for alternative methods to determine the pose of the ERF**

We tested additional methods other than the ones described in supplemental method 2 to characterize the relative pose of the TRF to the ERF, with the extra steps illustrated by Fig. S9. We outline processing steps relevant to segmenting the lens of the eye with a red box and steps relevant to segmenting the iris of the eye with a blue box.

1. First, we binarized the B-scan and generated a structural mask as detailed previously in supplemental methods 3.

2. In the binarized B-scan, we removed components spatially connected to the corneal top surface by removing the connected component closest to the outer surface of the eye to form a binarized image of the lens and part of the iris (Fig. S9a).
3. In the resulting binarized image, we kept connected regions with a bounding box that had an axial length greater than 40 µm and discarded the rest. We considered the largest remaining component as the lens (Fig. S9b) and fit a second-order polynomial to the outer surface of the lens between its left and right edges (Fig. S9c). In Fig. S9c, we illustrate the fitted outer surface with a red curve and the left and right edges of the lens with blue vertical lines.
4. To identify the boundaries of the iris, we designed a filter designed to amplify the signal of the iris in the vis-OCT images (Fig. S9d). The filter takes the signal amplitude of 3 axially adjacent pixels and subtracts the signal amplitude of the 8 pixels vertically adjacent to this region in the upwards and downwards direction. In vis-OCT, the iris appears as a thin surface owing to the attenuation of visible light by the pigmentation of the iris. We filtered the previewed B-scan using this filter (Fig. S9e).
5. The pixel of maximum intensity for each a-line not within the border of the lens was considered the axial position of the iris (Fig. S9f). In Fig. S9f, the blue vertical lines illustrate the border of the lens, and the red dots show the position of the iris.

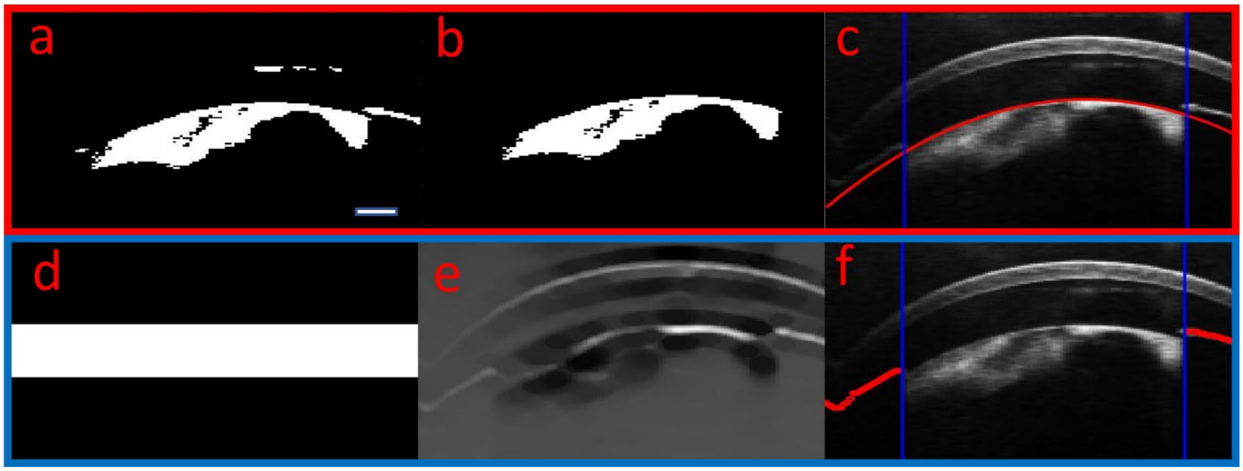


**Fig. S9. Real-time image processing to segment the lens and iris**. (a)-(c) Image processing related to determining the position of the iris. a) Binarized structural B-scan minus component touching the cornea. b) Post-processing of binarization to isolate lens. c) Quadratic fit of lens position in red lines and boundaries of the lens in blue lines. (d)-(f) Image processing related to determining the position of the iris. d) Filter designed to detect ridges, amplifying the signal of the iris. e) B-scan image filtered using filter in (d). f) Original B-scan with iris position overlayed in red and edge of lens given by blue lines. The scale bar is 200 µm.

**Supplemental Data 1: Evaluation of methods to identify the lateral origin of the ERF**

We tested the following image processing schemes for matching the transverse origin of the ERF and TRF, using the image processing steps detailed in supplemental methods 3 and 7:

1) **X center of mass:** Let I(x, z) be the signal amplitude of the B-scan at depth z and lateral position x. We define the x center of mass as

$$COM_x = \frac{\Sigma_z \Sigma_x I(x,z)x}{\Sigma_z \Sigma_x I(x,z)}. \qquad \text{Eq. S15}$$

We approximated the x center of mass as the origin of the ERF as the origin of the ERF along the axis that the B-scan was acquired.

2) **Cornea apex:** We considered the apex of the fitted cornea, found using the methodology in supplemental methods 3, to be the origin of the ERF along the axis where the B-scan was acquired (Fig. S10a).

3) **Lens Center:** After determining the boundaries of the lens, found in step 3 of supplemental methods 7, we took the midpoint between the lens boundaries to be the origin of the ERF (Fig. S10b).

To evaluate the accuracy of each method, we manually selected the lateral center of the central cornea as the ground truth and correlated the manually selected positions with the automatically calculated position. We computed the correlation of each method with the manually selected center and ordered the methods from the fewest to the most processing steps required (Fig. S10c). Overall, the cornea apex method had the greatest correlation.

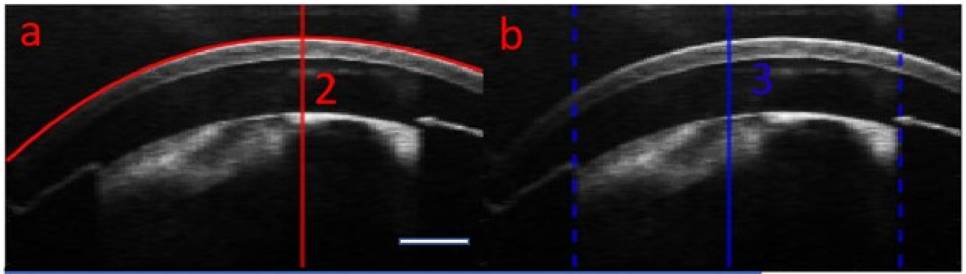


c

| Method | Correlation |
| --- | --- |
| 1. X Center of Mass | 0.917 |
| 2. Cornea Apex | 0.98 |
| 3. Lens Center | 0.684 |

Least steps → Most steps

**Fig S10. Schemes for real-time alignment of axes origin between robotic sample arm and eye**. (a) The top of the fit cornea is used to represent the lateral origin of the ERF along the X and Y axes. b) The center of the lens positions are used to represent the origin of the ERF along the X and Y axes. c) Correlation of different methods for determining eye origin, sorted from the least to most steps, compared to manual identification of origin. The scale bar is 200 µm.

**Supplemental Data 2: Evaluation of methods to identify the optical axis of the eye**

We tested the following methods for aligning the z-axis of the TRF and the Z-axis of the ERF. When the axes are misaligned, we note that the structures of the B-scan are tilted from the left and right sides of the image. Each of the following schemes was evaluated as a metric for assessing the relative orientation of the z-axis of the TRF and the Z-axis of the ERF.

1) **Y center of mass slope**: As detailed previously, we calculated the center of mass for the center B-scan in the axial direction as a function of transverse position x

$$COM(x) = \frac{\Sigma_z I(x,z)z}{\Sigma_z I(x,z)}. \quad \text{Eq. S16}$$

The slope of a line fit to $COM(x)$ (Fig. S11a) was used as a metric to assess how misaligned the two axes were.

2) **Filtered image y center of mass slope**: This method is identical to Y Center of Mass Slope except that the input image is the image filtered by the custom filter, found in step 4 of supplemental methods 7 (Fig. S11b). We show an example of the filtered image in Fig. S9e.

3) **Lens vector**: The average axial depth of the lens top border, found after step 3 of supplemental methods 7, on the left and right side of the lens was computed and the magnitude of the difference was taken (Fig. S11d).

4) **Lens line fit slope**: We fitted a slope to the top position of the binarized lens, found in step 3 of supplemental methods 3, from left to right.

5) **Iris vector**: The average axial depth of the iris, found after step 5 of supplemental methods 7, on the left and right side of the lens was computed and the magnitude of the difference was taken (Fig. S11d).

6) **Iris slope**: We calculated the slope of a line fit to the iris, found after step 5 of supplemental methods 7.

To evaluate the accuracy of each method, we manually drew a line parallel to the optical axis of the eye as the ground truth and measured the correlation of each method with the angle of the manually drawn line. We found that the y center of mass slope, iris vector, and iris slope had the highest correlations (Fig. S11e). Since the y center of mass slope involved the least computational steps, making it more efficient for real-time processing, we used the y center of mass slope for determining the optical axis of the eye.

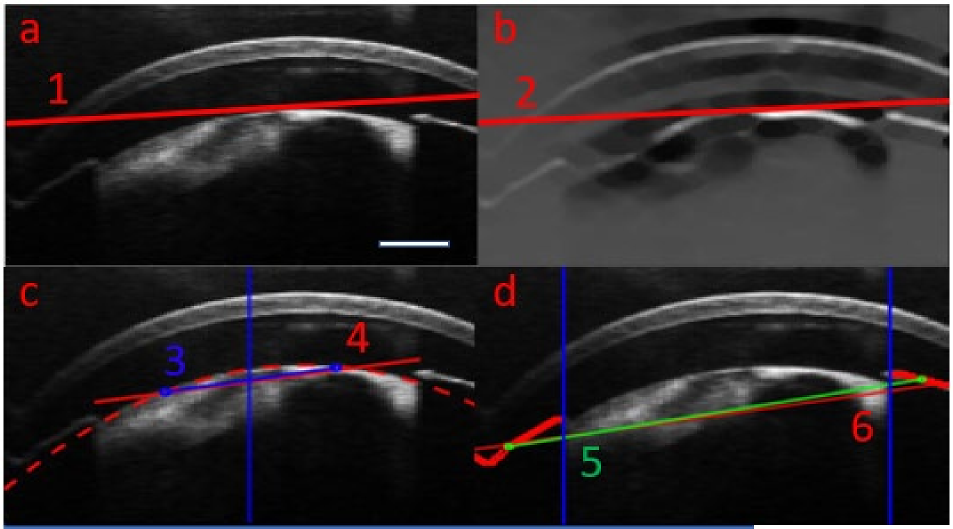


| e Method | Correlation |
|---|---|
| 1. Y COM slope | 0.967 |
| 2. Filtered image Y COM | 0.956 |
| 3. Lens vector | 0.71 |
| 4. Lens line fit slope | 0.779 |
| 5. Iris vector | 0.982 |
| 6. Iris slope | 0.981 |

Least steps → Most steps

**Fig S11. Schemes for real-time alignment of optical axes between TRF and ERF**. (a) (1) The slope of the vertical center of mass along the lateral position is used to estimate the alignment of the z-axis of the TRF and Z-axis of the ERF. b) (2) The slope of the vertical center of mass of the filtered image along the lateral position used to estimate the alignment of the axes. c) The difference in the center of the segmented lens along the left and right side of the lens (3) and slope fit along the lens top depth (4) used to estimate the alignment of the axes. d) The difference in average iris position to the left and right of the lens (5) and slope fit along iris position (6) used to estimate the alignment of the axes. e) Correlation of different methods for determining eye optical axes, sorted from the least to most steps, compared to manual identification of optical axes. The scale bar is 200 µm.

**Supplemental Data 3: Characterization of optical distortions**

To test how optical distortions influence our 3D reconstruction, we assessed the chromatic and radial distortions of our robotic vis-OCT. We theoretically calculated the beam size at 510, 560, and 610 nm at depths from 2mm above and below the OCT focal plane and 2mm to the left and right of the focal point using Zemax (Fig. S12a). We observed that the origins of each wavelength occupied the same spatial position.

To measure radial distortions, we imaged a grid distortion target (Thorlabs R1L3S3PR). The spacing between each grid was 100 μm. We imaged the distortion target using our vis-OCT (Fig. S12b) and measured the spacing between grid points as a function of distance away from the origin of the TRF (Fig. S12c). A linear fit of the number of lateral pixels between grid lines as a function of distance from the origin was $\#\, of\ pixels = 0.0005\ (\mu m\ from\ center) + 24.29$, with the slope not statistically different from zero with a p-value of 0.822. Thus, the lateral pixel size does not change substantially depending on the distance of the pixel from the center of the OCT FOV. Consequently, we found that radial distortion had minimal impact on the reconstructed volume. As the lateral dimension of the imaged grid distortion target was twice as large as what we used to acquire each volume of the outflow pathway and the number of pixels per grid line did not change as a function of distance from the TCP, correction of radial was not necessary for our robotic vis-OCT.

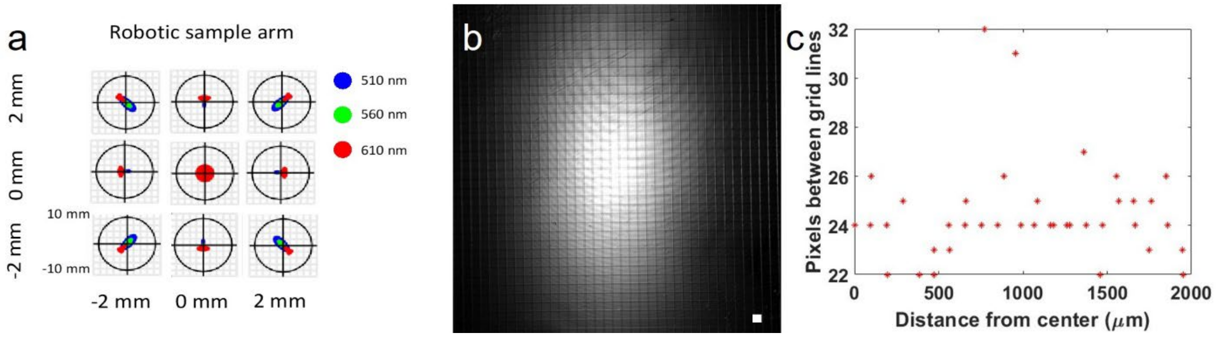


**Fig. S12. Characterization of optical aberrations.** a) Beam size at the focal plane at different wavelengths, depths, and positions. b) Image of distortion target grid, where the spacing between grid lines is 100 μm. c) Spacing between grid lines as a function of distance from the center of the OCT field of view.

**Supplemental Data 4: Validation of Resolution**

Using the radius of the airy disk definition, we calculated the theoretical lateral resolution of our robotic vis-OCT to be 9.4 µm. We validated our lateral resolution experimentally with a USAF51 target card. The smallest feature we could resolve on the target card was group 5, element 6 (Fig. S13a), which has a line width of 8.77 µm. The theoretical axial resolution of our vis-OCT is approximately 2 µm in air and 1.3 µm in tissue.

We validated the resolution of our vis-OCT by imaging a customized microfabricated polydimethylsiloxane phantom. The phantom consisted of cylindrical columns distributed in a uniformly spaced pattern. We characterized the structure of the phantom using a three-dimensional optical profiler (NexView, Zygo Corp). Each cylindrical column was 9.0 µm in height and 11.5 µm in diameter. The gap between columns was 8.5 µm (Fig. S13b). Using our vis-OCT, we imaged the our phantom (Fig. S13c). The columns and spacing in between the columns were visualized, indicating that our vis-OCT had sufficient resolution to discern the 8.5 µm gap between the columns. Measurement of the column diameter and space between columns was 11.5 and 8.5 µm using vis-OCT, in agreement with the optical profiler. To assess the height of the column, we examined a cross-sectional B-scan of the phantom (Fig. S13d). We measured the height of the column as 8.7 µm, slightly under the value recorded using the optical profiler.

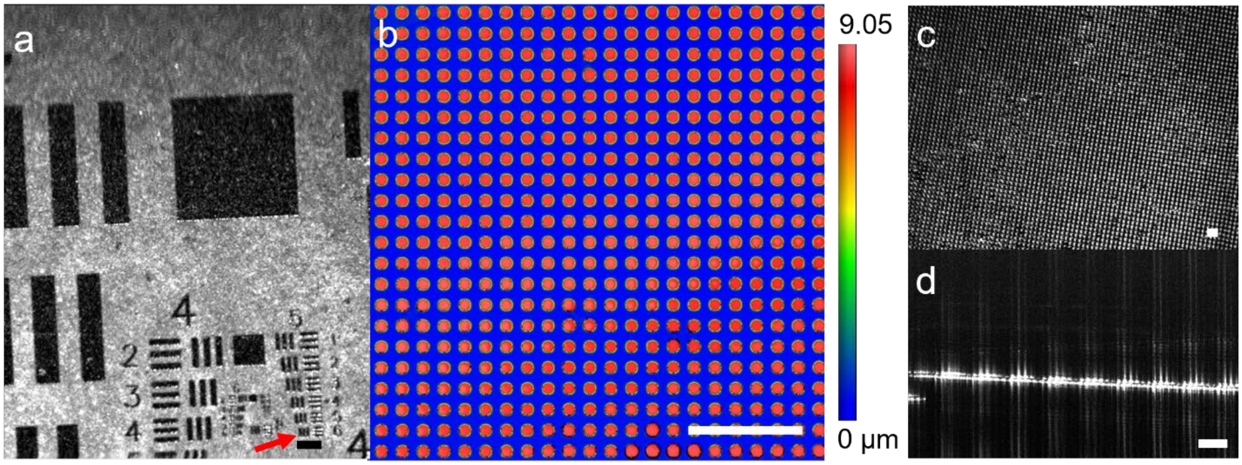


**Fig. S13. Characterization of lateral resolution of robotic vis-OCT.** a) En-face projection of a USAF51 target card. The red arrow points to group 5, element 6. The lines of group 5, element 6 were able to be separated. b) Customized microfabricated phantom characterized by a three-dimensional optical profiler. The red dots have a height of 9 µm and the blue background has a height of 0 µm. c) En-face projection of the microfabricated phantom imaged with vis-OCT. d) Cross-sectional B-scan of the microfabricated phantom used the determine the height of the columns. Scale bars are 100 µm.

**Supplemental Data 5: Repeatability of robotic AS-OCT positioning**

To monitor dynamic changes in structure, we desired that the positioning of the robot arm is repeatable over time and stable when held at a fixed position. To test the precision of our robot arm positioning, we attached a test cube to our robotic arm and tracked the position of that test cube with laser displacement sensors in 3 orthogonal directions (Fig. S14a). To measure repeatability, we moved the robotic arm to the same position 200 times. When commanded to return to the same position 200 times for five different poses, the standard deviations in measured positions in the x, y, and z directions were 2.4, 1.6, and 1.9 $\mu m$ respectively (Fig. S14b). The fluctuation in positions, defined as half the max range of positions, was 5.9, 4.0, and 5.2 $\mu m$ in the x, y, and z axis respectively (Fig. S14c). To measure stability, we fixed the robotic arm in place and took position measurements every 2 seconds for 400 seconds for five different poses. When held in place, the standard deviation of position in the x, y, and z directions were 1.6, 0.9, and 0.9 $\mu m$ (Fig. S14d) with a maximum fluctuation of 3.7, 2.0, and 2.3 $\mu m$ respectively (Fig. S14e).

Next, we tested the precision of our X and Y axes origin finding and optical axis determination methods for the ERF used in our data acquisition. To quantify the precision of our methods, we applied aligned our TRF with the coordinate system of a ROWE anterior segment phantom eye model. For testing the lateral origin finding, we displaced the phantom cornea a distance between -2mm to 2mm from the center of the phantom in increments of 0.2 mm. We subsequently let the TRF align its lateral origin with the phantom and took the recorded the internal robot arm position once the robot arm had achieved a stable position (Fig. S14f). Fig. S15g shows the a linear fit of final position of the robotic arm as a function of distance from the origin. The slope of the linear fit was not statistically significant different from zero although the fitted slope was 4.4 $\mu m/mm$. The precision of our centering algorithm, defined as the standard deviation in final measured positions, was 13.9 $\mu m$.

For testing the optical axis alignment method, we misaligned the phantom eye and TRF z-axes between -5 to 5 degrees in increments of 1 degree. We subsequently applied the method to correct for the misaligned axes and recorded final orientation once the robot arm had achieved a stable position (Fig. S14h). Fig. S14i shows a linear fit of final angular misalignment, defined as a difference in orientation from the starting aligned pose, as a function of initial angular misalignment. The slope of the linear fit was not statistically significant different from zero although the fitted slope was 0.026 degrees change in final angular misalignment per 1 degree change in initial angular misalignment. The precision of our z-axes alignment algorithm, defined as the standard deviation of final angular misalignment, was 0.28 degrees.

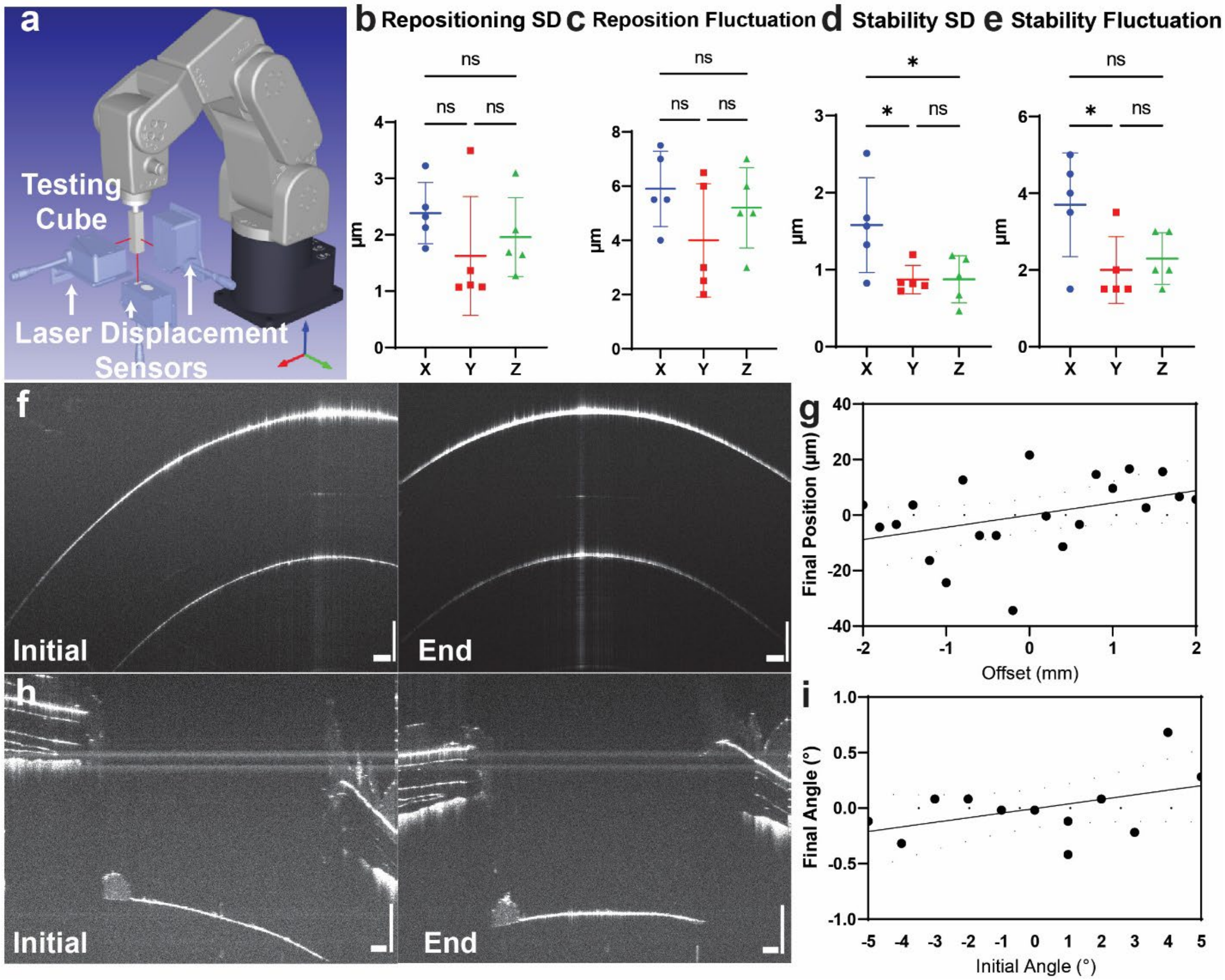


**Fig. S14. Precision of robotic AS-OCT system and control schematics.** a) Experimental setup for testing robotic repositioning and stability. b-c) Standard deviation and fluctuation (range of recorded positions) of robotic arm test cube position after repositioning cube. d-e) Standard deviation and fluctuation of test cube position when held in place. f) Centering algorithm applied to the phantom cornea. g) Final recorded position of the robotic arm as a function of offset from the correct center position. h) Tilting algorithm applied to phantom lens and iris. i) Final recorded position of the robotic arm as a function of offset from the correct starting angle.

**Supplemental Data 6: 3D reconstruction results in less biased measurements of eye radius**

We measured the radius of the mouse eye using our fully reconstructed digital twin of the eye and compared the measured value with that which would be measured using only cross-sectional images. To measure the radius of the eye, we fit a sphere to point cloud of eye outer surface (Fig. S15a). For an example BALB/c mouse, we measured the radius of curvature to be 1.7 mm. Next, we measured how much the radius of curvature when only examining individual cross sections by taking cross-sectional planes of the full point cloud (Fig. S15b). A circle was fit to each cross-sectional plane (Fig. S15c). For cross-sections passing through the center of the eye but orientation rotated at different angles around the eye, the measured radius of curvature was 1.7 ± 0.2 mm (Fig. S15d). Overall, utilizing the full 3D data avoids variation in measured radius that would occur if looking at specific planes. Next, we moved a cross-section oriented along the y-direction of the ERF away from the center of the eye. The measured radius of curvature decreased the farther the cross-section was from the center (Fig. S15e).

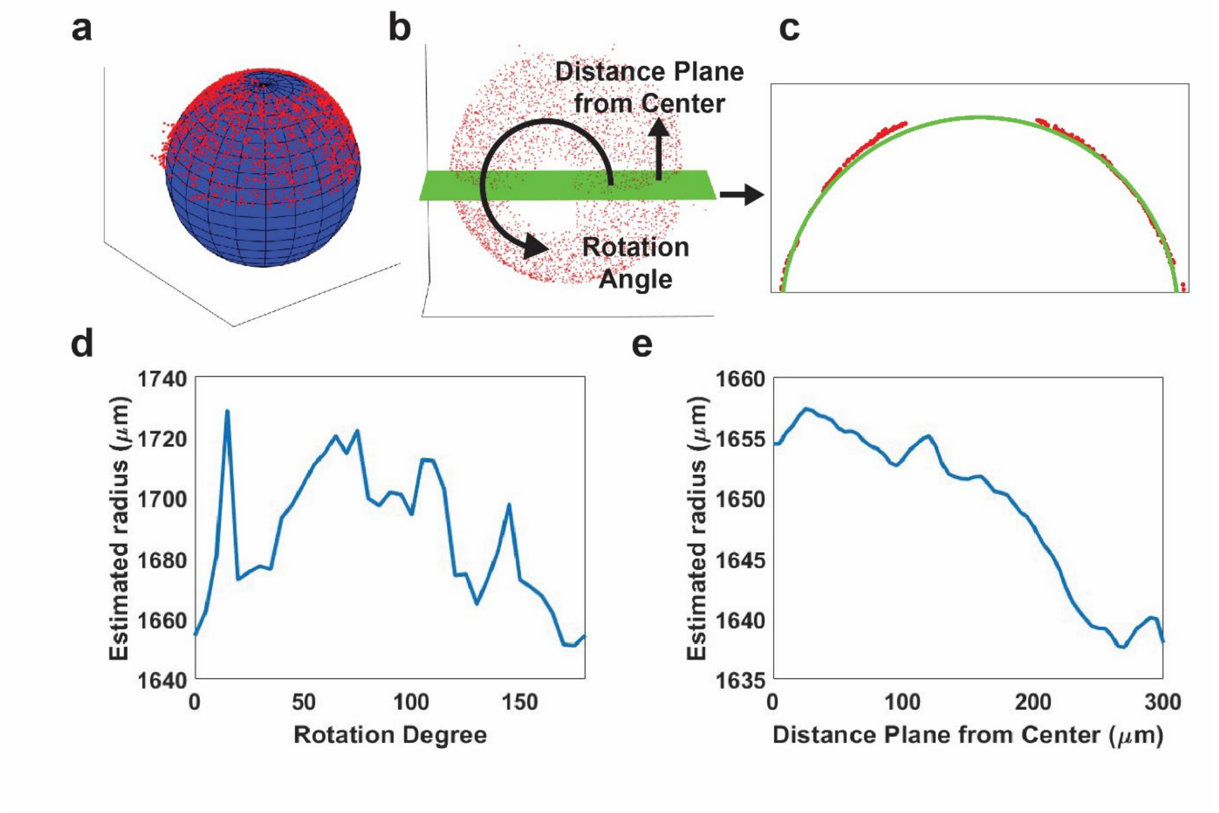


**Fig. S15. Full volumetric reconstruction for determining eye radius.** a) The radius of the eye was determined by fitting a sphere to the point cloud representing the outer surface of the eye (red points). b) The radius of the eye was also estimated by fitting a circle to the outer surface of cross-sections of the eye. The cross-section was rotated and translated to assess how the measured radius varies with the choice of cross-section. c) Fitted circle to cross-section of eye (red points) was used to estimate the radius of curvature. d) Estimated radius of curvature as cross-section is rotated about the center of the eye. e) Estimated radius of curvature as cross-section is translated away from the center of the eye.

**Supplemental Data 7: Repeatable Positioning Enables Analysis of Dynamic Changes**

A primary advantage of *in vivo* imaging over histology is the capability to assess functional properties of tissue. To assess the dynamic response of SC to external modulation, we imaged the full conventional outflow pathway in a C57BL/6 mouse before and 15 minutes after 1% pilocarpine administration. Pilocarpine is expected to increase SC size by constructing the pupil and pulling on the zonule fibers connecting the TM to the lens. The pull of the TM outwards expands SC size[10,11].

Consistent with pilocarpines mechanism of action, the pupil constricted (Fig. S16 a-b)[12]. The outer edges of the lens is easily observed in the OCTA projection before pilocarpine adminsitation and not 15 minutes after administration. Increased OCTA signal was observed in the nasal-superior region and decreased signal was observed in the temporal-inferior region. The green dotted region shows an area with decreased OCTA signal after administration and the red dotted area shows a region where a new vessel branch is observed. These patterns were consistent with regional changes in SC volume following pilocarpine administration, where SC volume per degree of eye increased the most in the nasal-superior region and least in the temporal-inferior region (Fig. S16c). Volume per degree of the eye was found by computing the number of voxels within one azimuthal degree of the final reconstructed volume. Volume values were averaged over a region of 20 degrees for stable readings. For this mouse, we found that volume increase per degree of the eye following pilocarpine administration was 40.5 ± 30.8% and ranged from 3.2 to 116.7%. Overall, the total volume increased by 15.5% in the inferior quadrant, 84.4% in the nasal quadrant, 39.7% in the superior quadrant, and 26.7% in the temporal quadrant. Just as with SC size, we found a segmental variation in volume changes following pilocarpine administration.

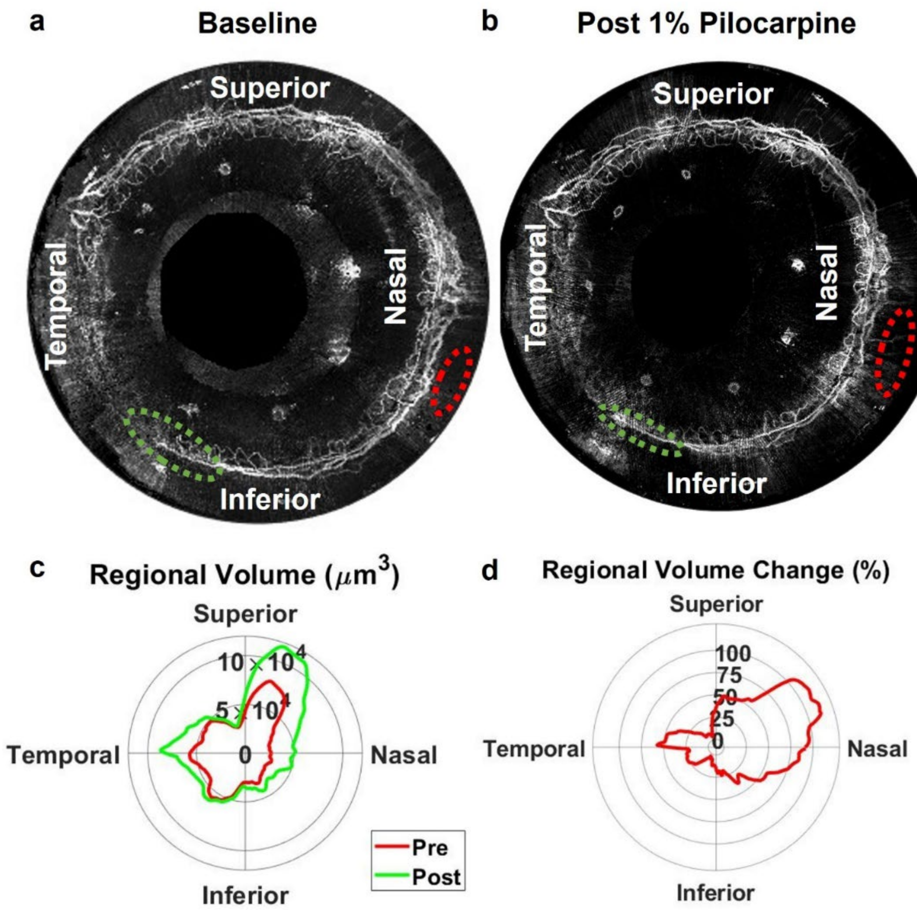


**Fig. S16. Segmental dynamic changes in SC morphology captured after pilocarpine administration.** a)-b). Projection of angiogram prior to and 15 minutes after 1% pilocarpine administration. Decreased OCTA signal in the inferior region and increased signal in the nasal region were observed. The dotted green circle illustrates a region where OCTA signal decreased after pilocarpine administration and the dotted red circle illustrates a region where OCTA signal increased after pilocarpine administration, with the ability to visualize another vessel branch. Pupil size decreased following pilocarpine administration. c) Segmental volume within 1 degree of eye per quadrant prior to and after pilocarpine administration. Volume values averaged within 20 degrees. d) Volume increase as a percentage of the original volume following pilocarpine administration.
